# SCKAnalyzer: A Web Tool for Single-Cycle Kinetics Analysis and Simulation

**DOI:** 10.64898/2026.10.07.756993

**Authors:** Florian Malard, Jean-Michel Blanc, Eric Roubin, Thomas Schäfer, Carmelo Di Primo

**Author notes:** Deceased, August 31, 2026 (Eric Roubin). Corresponding authors. Univ. Bordeaux, CNRS, INSERM, ARNA, UMR 5320, U1212, IECB, F-33000 Bordeaux, France *E-mail addresses:* (F. Malard), (C. Di Primo).

## Abstract

Surface plasmon resonance (SPR) and biolayer interferometry (BLI) are well-established label-free analytical technologies used to determine biomolecular interaction rates (*k*_a_, *k*_d_) and equilibrium constants (*K*_D_). While assays historically rely on multi-cycle kinetics (MCK), this approach requires intermediate surface regeneration, risking sensor degradation and increasing reagent consumption. Single-cycle kinetics (SCK) avoids regeneration by sequentially injecting increasing analyte concentrations. However, implementing SCK workflows on biosensor platforms like the Octet^®^ R8 BLI instrument (Sartorius) remains a challenge due to the lack of built-in or online data analysis tools.

To address this, we introduce SCKAnalyzer, a free, user-friendly web application designed for processing and fitting SCK datasets. SCKAnalyzer accepts column-based text files and native raw BLI data (.frd format) from Sartorius instruments, performing data pre-processing and non-linear fitting to a 1:1 interaction model. Additionally, it offers advanced simulation capabilities that allow variable association and dissociation durations across sequential injection steps, a feature absent in existing platforms such as Biacore™ Simul8.

The performance of SCKAnalyzer was experimentally validated using a 3-step SCK method (SCK3) for DNA/RNA duplex formation on a Biacore T200, as well as 4-step (SCK4) and 5-step (SCK5) methods for DNA/DNA duplex formation on an Octet^®^ R8 BLI instrument. Binding parameters (*k*_a_, *k*_d_, *K*_D_) derived from SCKAnalyzer showed excellent agreement with values from established proprietary softwares such as Biacore T200 Evaluation Software v3.2.1 and BIAevaluation v4.1.1. SCKAnalyzer provides a practical alternative to MCK, reducing probe consumption and extending SCK utility across biosensor platforms. SCKAnalyzer is available at sck.iecb.u-bordeaux.fr (source code: github.com/synthaze/sckanalyzer).

## 1. Introduction

In the 1990s, surface plasmon resonance (SPR) emerged as a powerful label-free technology for monitoring biomolecular interactions in real time [1–4]. By measuring mass changes resulting from complex formation at a solid/liquid interface, SPR-based instruments as those pioneered by Biacore (now Cytiva, Uppsala, Sweden) quickly became the standard in both academic and industrial laboratories for determining the association and dissociation rate constants, *k*_a_ and *k*_d_, respectively, as well as the dissociation equilibrium constant, *K*_D_ [5–8].

For decades, SPR faced no competition and is still viewed as the gold-standard for characterizing the interaction kinetics involving biomolecules. In recent years instruments based on other physical principles and with different specifications have successfully entered the market. Among them, biolayer interferometry (BLI)-based biosensors quickly met expectations by enabling experiments to be conducted faster and at lower costs than SPR in most settings [9–11]. While SPR monitors mass changes in real time within a microfluidic system, on Biacore instruments, using sequential injections of the analytes, BLI monitors changes in optical thickness at the surface of a disposable tip immersed in the samples using light interference, in a parallel format, which greatly reduces experiment duration. Ultimately both platforms yield the same binding parameters, *k*_a_, *k*_d_ and *K*_D_. Beyond SPR and BLI, other commercially available technologies for determining these parameters have their own advantages and disadvantages. Therefore, no single technique can fully supplant the others. For instance, integrating both BLI and SPR was recently shown to be beneficial [12].

Historically, SPR assays have relied on a multi-binding cycle approach that requires the functionalized sensor chip to be regenerated between each injection of the analyte at increasing concentrations [3,7,9,13–15]. The method remains the most widely used across all available instruments, regardless of the measurement principle. While highly effective, it presents challenges when dealing with unstable molecules, regeneration-sensitive biosensor surfaces, or expensive reagents. To circumvent these limitations, Karlsson et al. introduced a method in 2006 known as single-cycle kinetics (SCK), also referred to as kinetic titration, and explicitly formalized the term multi-cycle kinetics (MCK) to refer to the traditional multi-regeneration approach [16]. In an SCK assay, sequential injections of the analyte at increasing concentrations are flowed over the functionalized biosensor, without regeneration steps between sample injections, which makes SCK faster than MCK and minimizes surface degradation. Ultimately SCK reduces costs.

Despite the advantages of SCK and its availability on Biacore instruments for two decades, MCK is still preferred over SCK. This is further illustrated by BLI-based systems. Even though performing SCK assays on instruments such as Octet^®^ (Sartorius) and Gator (Gator Bio) is straightforward, their native software packages do not include built-in tools for SCK kinetic evaluation. In 2017 Kamat and Rafique successfully performed SCK on an Octet HTX (Sartorius) fitting the data with the Biacore’s original BIAevaluation v4.1.1 software [17]. This commercial tool from Biacore (Cytiva) is not designed for this purpose but it allows users familiar enough with math and basic coding to write the equations needed for SCK analysis, in a Biacore-based programming language. Despite being over two decades old, this proprietary software remains a powerful tool, which can be used also for modelling equations that describe complex binding reactions and experimental setups [18,19]. However, it is clearly not accessible to all users. In 2019, Krämer et al. introduced *Anabel*, an online tool that initially did not include SCK analysis [20]. While its web-based version has since been discontinued, the tool can still be run locally using R and now includes SCK analysis. More recently, additional tools capable of SCK analysis have been developed, including the Mathematica package TitrationAnalysis [21] and a Python Jupyter notebook [22]. Conversely, other tools such as Scrubber [7] and TraceDrawer (commercial), KinGenie (web-based and free) or KinetiHub (web-based and partially free) continue to support only MCK analysis.

Given the effectiveness of the SCK method and the lack of web-based analysis options, we introduce SCKAnalyzer, a free and user-friendly online tool designed for SCK data analysis. Intended primarily to address our need to analyze results obtained by SCK using an Octet^®^ R8 BLI instrument (Sartorius), SCKAnalyzer can also be used with data from other instruments. The online tool allows users to upload raw data (.frd files) from the Octet^®^ R8, as well as .csv or .dat/.txt files exported from it or other instruments, apply pre-processing steps and fit the sensorgrams to a 1:1 interaction model to determine *k*_a_, *k*_d_ and *K*_D_. SCKAnalyzer, unlike Biacore™ Simul8 (Cytiva), also enables SCK simulations with variable association and dissociation durations prior to the final dissociation step, a crucial feature for any system that support adjustable step durations, such as BLI instruments. To validate SCKAnalyzer, we first monitored DNA/RNA duplex formation via SCK using three analyte concentrations (SK3) on a Biacore T200. We first processed the data with the instrument’s dedicated software Biacore T200 Evaluation v3.2.1, and then analyzed them with Biacore BIAevaluation v4.1.1 and with our platform. To further validate our tool, we monitored DNA/DNA duplex formation by SCK on an Octet^®^ R8 BLI instrument under two different SCK conditions, four (SCK4) and five (SCK5) analyte concentrations, and different kinetic regimes. For a direct comparison, we analyzed the sensorgrams using SCKAnalyzer and BIAevaluation v4.1.1.

The results demonstrate that SCKanalyzer, a free and user-friendly web-based application, enables reliable, instrument-independent kinetic analysis of sensorgrams obtained by the SCK method. Used on a BLI instrument, SCK reduces the number of probes required per experiment. Although this increases assay duration relative to parallel MCK, the key technical feature that popularized BLI instruments, the added time is not a major drawback in many cases. Overall, combining SCK with SCKAnalyzer offers a practical alternative to conventional MCK, extending its utility beyond BLI instruments to any platform capable of monitoring kinetics through titration assays.

## 2. Material and Methods

### 2.1. Webserver implementation

SCKAnalyzer is implemented as a stateless client-server web application. The backend is a REST API written in Python 3.11 using the FastAPI framework (v0.115) and served over Uvicorn (v0.30), an ASGI server. Numerical routines rely on NumPy (v2.0) [23], pandas (v2.2), and SciPy (v1.14) [24]. Uploaded sensorgrams, either generic delimited text files or Octet BLI XML (.frd) exports, are parsed and processed entirely in memory for the duration of a single request. No database or server-side session state is used, and no uploaded data or fitted results are persisted after the response is returned. Parameter estimation is performed with the bounded non-linear least-squares solver provided by SciPy (*scipy.optimize.least_squares*) [24]. The client is a single-page application written in TypeScript with React (v18), bundled with Vite (v5), and uses Plotly.js for interactive sensorgram, residual, and overlay plots. It communicates with the backend exclusively through a small set of JSON/multipart REST endpoints (file parsing, single-replicate fitting, global multi-replicate fitting, and a health check). The application is deployed as a single-host stack on a Debian-based GNU/Linux distribution, with the backend managed as a systemd service behind an nginx reverse proxy. A live instance is hosted at sck.iecb.u-bordeaux.fr. The source code is released under the GNU General Public License v3.0 and is publicly available at github.com/synthaze/sckanalyzer. Claude Code (Anthropic), using the Claude Opus 4.8 model, was used to assist in the development of the application.

### 2.2. Kinetic modeling and curve fitting

Sensorgrams are modeled with a pseudo-first-order 1:1 Langmuir binding scheme [3,25],

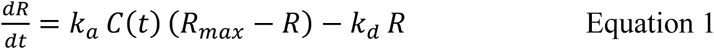

Where *R* is the biosensor response, *C(t)* the analyte concentration along the injection schedule, *k*_a_ and *k*_d_ the association and dissociation rate constants, respectively, and *R_max_* the maximal binding capacity. Because *C(t)* is piecewise constant across the injection and wash windows of a single-cycle run, the equation is solved in closed form on each segment: with

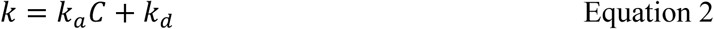

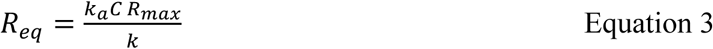

(*R_eq_* = 0 when *C* = 0), the response evolves as

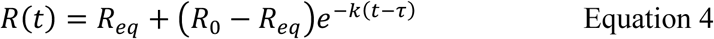

where *τ* is the start time of that segment and *R*_0_ the response carried over from the preceding one, enforcing continuity across the trace. A constant bulk (refractive-index) offset may additionally be applied within each injection window. The equilibrium dissociation constant is reported as

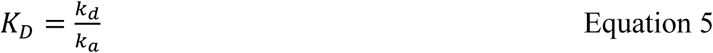

Parameters (log_10_*k_a_*, log_10_*k_d_*, *R_max_*, and any per-injection bulk offsets) are estimated by non-linear least-squares, minimizing the residuals between predicted and observed response over the entire multi-injection trace at once rather than segment by segment. Rate constants are optimized in log_10_ space to enforce positivity and to span their multi-order-of-magnitude range, and a user-selectable robust loss (linear, soft-L1, or Huber) down-weights outlying residuals such as injection spikes. Traces may first be corrected by subtraction of a reference channel and/or a referenced blank channel. User-defined time windows (e.g., bubbles or spikes) are excluded from the objective function and from all reported metrics, but remain visible in the displayed trace and residual plot. For experiments with several replicates, a global fitting mode shares *k*_a_ and *k*_d_ across replicates, optionally together with *R*_max_ and the bulk offsets, minimizing their concatenated residuals jointly.

For global fitting conditions, parameter uncertainties are primarily reported as asymptotic standard errors obtained from the Jacobian *J* of the residuals at the optimum:

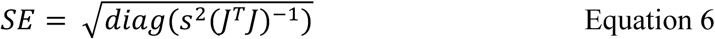

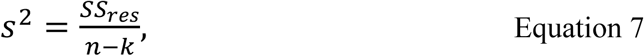

where *n* is the number of fitted points and *k* the number of free parameters. In per-replicate fitting, mean parameters with standard deviations are reported in addition to per-replicate parameters with standard errors. Overall fit quality is summarized by the root-mean-square error, the coefficient of determination, and the reduced chi-square statistic, computed on the same points as the standard errors above.

### 2.3. Surface plasmon resonance experiments

SPR experiments were carried out at 25 °C using a Biacore T200 instrument (Cytiva, Uppsala, Sweden). 70 resonance units (RU) of a modified 5’-biotin-TEG-DNA oligonucleotide with the sequence _5’_TCACAAGTTAGGGTCTCAGGGA_3’_ (Eurofins Genomics, Nantes, France), prepared at 50 nM in PBS buffer (Invitrogen, Thermo Fisher Scientific, Villebon-sur-Yvette, France) containing 0.05% Tween-20 (running buffer), were immobilized by streptavidin-biotin coupling onto SAHC200m sensor chip (Xantec Bioanalytics, Düsseldorf, Germany). A flow cell was left blank for double refencing of the sensorgrams. A complementary RNA strand, _5’_UCCCUGAGACCCUAACUUGUGA_3’_, was used as analyte. It was synthetized in the laboratory using a K&A H8 DNA/RNA synthesizer (Sierra BioSystems, Schaafheim, Germany) and purified by electrophoresis on denaturing polyacrylamide gels. The RNA oligonucleotide, prepared in the running buffer at three concentrations (5, 50 and 500 nM), was injected for 45 s over the functionalized surface at 25 µL/min flow rate using an SCK method consisting of three sequential injections (SCK3), performed in duplicate. Regeneration of the surface was achieved with a 30 s pulse of 20 mM NaOH prepared in milli-Q water. Sensorgrams were fitted to a 1:1 Langmuir model of interaction to determine the association rate constant, *k*_a_, and the dissociation rate constant, *k*_d_. The dissociation equilibrium constant, *K*_D_, was calculated as *k*_d_/*k*_a_. Curve fitting was first performed using Biacore T200 Evaluation v3.2.1 software. Then the double referenced sensorgrams were exported as .txt files and uploaded to the Biacore’s original BIAevalution v4.1.1 software and to SCKAnalyzer.

### 2.4. Biolayer interferometry experiments

BLI assays were performed using an Octet^®^ R8 instrument (Sartorius France, Dourdan, France) with SAX streptavidin probes (Sartorius). For these experiments a modified 5’-biotin-TEG-DNA oligonucleotide with the sequence _5’_GGACTTCGGA_3’_ was used as ligand, and its complementary DNA strand _5’_TCCTGAAGTCC_3’_ (Eurofins Genomics) as analyte. Two SAX probes (Sartorius) were positioned in column. One probe was functionalized with 0.2 nm of biotinylated target (target probe), prepared at 25 nM in HBS-EP+ (probe immersion for 180 s). The other one was left blank (blank probe) for double referencing of the sensorgrams. Both probes were then immersed for 3 min in a 1.2 µM solution of biotin prepared in HBS-EP+ to block any unoccupied streptavidin sites. SCK assays with four analyte concentrations (SCK4) were performed at 25 °C in 10 mM sodium phosphate buffer, pH 7, containing 50 mM NaCl and 0.05% Tween-20, while those with five analyte concentrations (SCK5) were conducted at 23 °C in HBS-EP+ running buffer (Cytiva, containing 150 mM NaCl). For SCK4, the target and blank probes positioned in column were immersed sequentially for 60 s (70 s dissociation phase between injections) in two rows of wells filled with the complementary strand at increasing concentrations of 62.5, 250, 1000, and 4000 nM. For SCK5, the probes were immersed sequentially for 30 s (60 s dissociation phase between injections) in wells with the complementary strand prepared at 14.8, 44.4, 133.3, 400, and 1200 nM. SCK4 and SCK5 assays were performed in duplicate. For both methods, one experiment was performed using buffer-only wells to allow double-referencing of the sensorgrams. The BLI data were either exported as .dat files (txt files) using Discovery software and uploaded to BIAevaluation v4.1.1 or uploaded directly as .frd files to SCKAnalyzer. Sensorgrams were double referenced prior to fitting with a 1:1 Langmuir model of interaction to determine *k*_a_, *k*_d_ and *K*_D_.

## 3. Results and Discussion

### 3.1. Overview of the analysis workflow

SCKAnalyzer implements the single-cycle kinetics (SCK) analysis pipeline as a sequential, browser-based workflow, introduced by a video tutorial on the landing tab and described in Figure 1. The next three tabs each handle one analysis stage, with data passed forward automatically: corrections made during data preparation carry through step definition into the fit, so the curve inspected at every stage is the one ultimately fitted. A final tab simulates SCK experiments to help design conditions for measuring biomolecular interactions.

**Figure 1:**
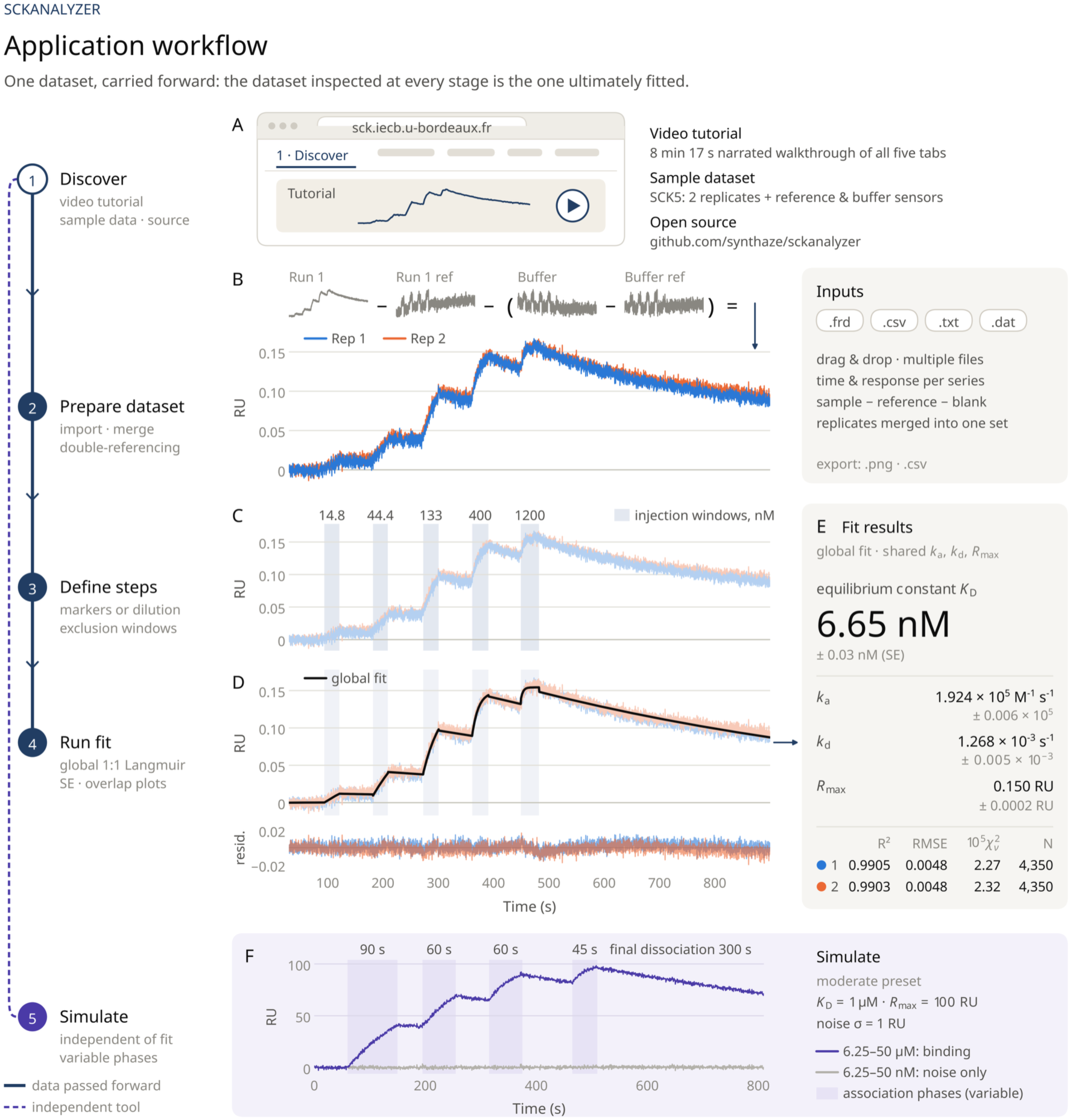
SCKAnalyzer application workflow. The browser-based analysis proceeds sequentially through the application tabs (left rail). Data pass automatically from the import tab to the fitting tab (solid line), and the simulator runs independently (dashed line). (A) Landing tab with video tutorial, sample dataset and source code. (B) Data import (.frd, .csv, .txt, .dat) with double referencing: each replicate is corrected by its reference channel and a referenced blank run. This is shown for one replicate on raw Octet BLI traces, with the corrected sensorgrams of both replicates below. (C) Injection steps: shaded association windows with their analyte concentrations. (D) Global 1:1 Langmuir fit (black; kₐ, k_d and R_max shared across replicates) with residuals. In (C–D), earlier layers are muted to highlight each new element. (E) Fitted kinetic and affinity constants with asymptotic standard errors, and per-replicate fit quality (R^2^, RMSE, reduced χ^2^, number of points). (F) Simulated moderate-affinity interaction with noise, variable phase durations and a final dissociation. Nanomolar injections show no detectable binding (gray), whereas micromolar injections reach saturation (violet).

### 3.2. Data import and reference subtraction

Users upload one or more sensorgram files, either generic .csv, .txt or .dat tables or native Octet^®^ BLI .frd files, by drag-and-drop or file selection. Multiple files and multiple replicate series per file are supported. For each replicate, users assign time and response columns and can subtract a reference channel from another uploaded replicate, with an optional "double-reference" step that subtracts a second, already-referenced blank run. Replicates spread across files can be merged into a single working set. The corrected trace appears immediately as an interactive sensorgram, and both the plot and its data table can be exported as .png or .csv files.

### 3.3. Injection step and exclusion window definition

The second stage defines the time window of each analyte injection, either by placing paired start and end markers on the sensorgram or by generating a full step table from a few serial-dilution parameters. For convenience, we have chosen to retain the term “injection” even though BLI instruments do not strictly inject samples, as the software can be used with other sample-injecting instrument. The step table remains editable either way. Users can also define exclusion windows to remove artifacts such as injection spikes or air bubbles before fitting. Injection steps and exclusion windows are overlaid on the sensorgram for visual verification.

### 3.4. Langmuir kinetic fitting and quality assessment

The third stage configures and runs the fit on the reference- and blank-subtracted trace from the first stage, using the injection steps and exclusion windows just defined. All fits use a 1:1 Langmuir binding model, deliberately, to prevent over-fitting: complex models almost always fit better numerically without reflecting the true biological interaction, often merely capturing experimental artifacts. Replicates can be fitted independently or globally, sharing the rate constants *k*_a_ and *k*_d_ and optionally *R*_max_. Options include per-injection bulk-offset fitting, shareable in global mode, an optional baseline correction applied before fitting, and a robust soft-L1, linear or Huber loss for the nonlinear least-squares optimizer. Each of *k*_a_, *k*_d_ and *R*_max_ can be toggled and bounded. Fitted curves and residuals are shown per replicate, or overlaid across replicates in global mode, alongside association and dissociation overlap plots that align each injection segment on a common time origin and report its RMSE. The estimates of *k*_a_, *k*_d_, *K*_D_ and *R*_max_ are reported with asymptotic standard errors from the Jacobian-based covariance estimate, optional residual-resampling bootstrap 95 % confidence intervals, and a replicate-wise mean ± SD summary when appropriate. Fit quality is summarized by RMSE, R^2^, reduced χ^2^ and the number of points used. All plots, parameter tables and overlap data can be exported individually as .png or .csv files, or together as a single .zip archive.

### 3.5. SCK experiment simulation

A separate tab generates synthetic single-cycle sensorgrams from user-specified kinetic parameters and injection schemes, independently of the fitting workflow, to explore expected instrument responses or produce test data. It builds the dilution series automatically, with independently configurable inter-injection and final dissociation windows and an optional linear drift term. Notably, SCKAnalyzer allows association and dissociation phases of variable duration before the final dissociation step, a feature missing from Cytiva’s Biacore™ Simul8 online simulation tool (https://apps.cytivalifesciences.com/spr/). This is particularly relevant for technologies such as BLI and any other instruments that support variable association and dissociation times across sequential injections.

### 3.6. Analyzing SPR and BLI data with SCKAnalyzer: application and performance evaluation

Although there are now many techniques available for measuring interaction kinetics, SPR remains the gold standard. Therefore, to evaluate how SCKAnalyzer performed we used first data obtained with a Biacore T200 with an SCK method consisting of three sequential injections of the analyte. Results from the global analysis of the two replicates are presented in Figure 2A and the DNA/RNA duplex formation parameters are listed in Table 1 (see Figure S1 for the analysis by replicate, together with the global analysis). *K*_D_ is in the sub-nanomolar range, 0.26 nM. The interaction is characterized by a slow dissociation phase, consistent with the 22-mer length of the DNA/RNA hybrid and the ionic strength of the PBS buffer. These double-referenced data were then exported as .txt files and uploaded to BIAevaluation v4.1.1. Although this legacy Biacore software lacks the equations for SCK data analysis, custom equations can be written in the “Edit Models” menu as described previously (ref Karlsson and Di Primo). The results from the analysis are presented in Figure 2B and the corresponding parameters are listed in Table 1 (see Figure S2 for the analysis by replicate, together with the global analysis). As anticipated, the parameters obtained from BIAevaluation v4.1.1 closely aligned with those from Biacore T200 v3.2.1, two well-established analysis tools. The .txt files were then uploaded to SCKAnalyzer. The results are presented in Figure 2C and the corresponding parameters are listed in Table 1 (see Figure S3 for the analysis by replicate, Figure S4 and Figure S5 for the parameters, together with those from the global analysis). The results demonstrate that SCKAnalyzer performs very well by successfully recovering the results obtained with Biacore tools, thereby validating the consistency across the different software packages.

**Figure 2:**
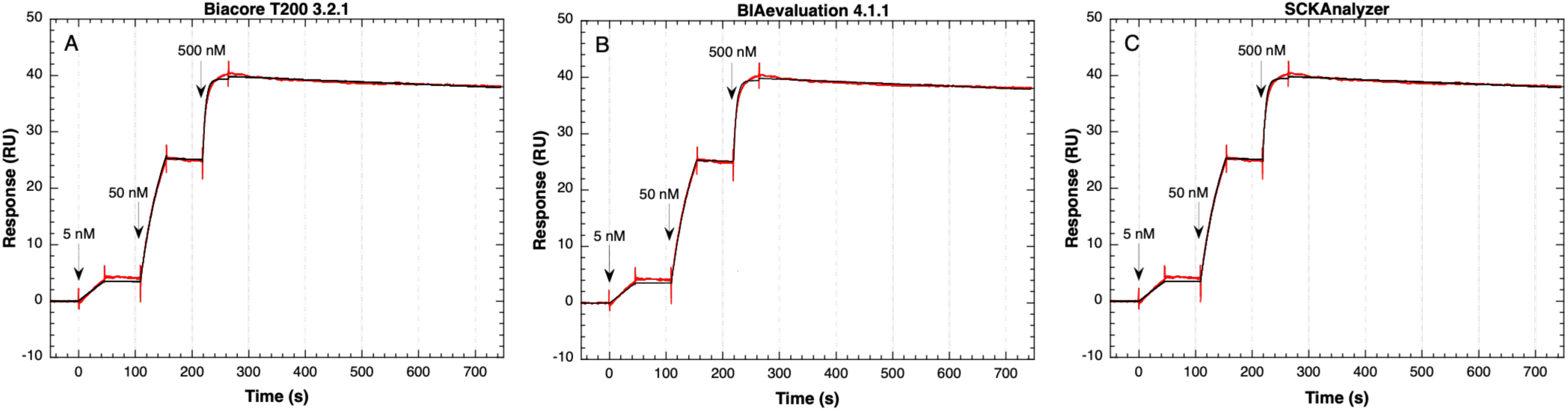
Sensorgrams of the DNA/RNA duplex, analyzed across three different computational tools. The red and black lines represent the experimental data and fit to a 1:1 model of interaction, respectively. Arrows indicate injection start times.

**Table 1.** Binding parameters obtained from various analysis software using the SCK method, consisting of three analyte injections (SCK3) for the SPR experiments, and four (SCK4) and five (SCK5) for the BLI experiments.

| | | $k_a$<br>$\pm \text{SE}^a (\text{M}^{-1}\text{s}^{-1})$ | $k_d$<br>$\pm \text{SE} (\text{s}^{-1})$ | $K_D$<br>(nM)<sup>b</sup> | $R_{\text{max}}$<br>$\pm \text{SE} (\text{RU})$ | Chi<sup>2</sup> |
| --- | --- | --- | --- | --- | --- | --- |
| SCK3 | Biacore T200<br>v3.2.1 | $4.037 \times 10^5$<br>$\pm 2.6 \times 10^2$ | $1.053 \times 10^{-4}$<br>$\pm 6.4 \times 10^{-7}$ | 0.2608 | 39.85<br>$\pm 7 \times 10^{-3}$ | 0.115 |
| | BIAevaluation<br>v4.1.1 | $4.10 \times 10^5$<br>$\pm 2.65 \times 10^2$ | $1.05 \times 10^{-4}$<br>$\pm 6.38 \times 10^{-7}$ | 0.256 | 39.8<br>$\pm 7.06 \times 10^{-3}$ | 0.116 |
| | SCKAnalyzer | $4.09 \times 10^5$<br>$\pm 2.81 \times 10^2$ | $1.034 \times 10^{-4}$<br>$\pm 6.659 \times 10^{-7}$ | 0.253<br>$\pm 1.637 \times 10^{-3}$ | 39.82<br>$\pm 7.45 \times 10^{-3}$ | 0.116<sup>c</sup> |
| SCK4 | SCKAnalyzer | $5.167 \times 10^4$<br>$\pm 4.889 \times 10^2$ | $1.0647 \times 10^{-2}$<br>$\pm 5.610 \times 10^{-5}$ | 206.04<br>$\pm 2.231$ | 0.09545<br>$\pm 3.39 \times 10^{-4}$ | 2.93 x<br>10<sup>-5c</sup> |
| | BIAevaluation<br>V4.1.1 | $5.20 \times 10^4$<br>$\pm 9.79 \times 10^2$ | $1.06 \times 10^{-2}$<br>$\pm 1.87 \times 10^{-4}$ | 203.8 | 0.0954<br>$\pm 3.9 \times 10^{-4}$ | 2.97 x<br>10<sup>-5</sup> |
| SCK5 | SCKAnalyzer | $1.924 \times 10^5$<br>$\pm 6.485 \times 10^2$ | $1.279 \times 10^{-3}$<br>$\pm 4.984 \times 10^{-6}$ | 6.651b<br>$\pm 3.426 \times 10^{-2}$ | 0.150<br>$\pm 1.53 \times 10^{-4}$ | 2.31 x<br>10<sup>-5c</sup> |
| | BIAevaluation<br>V4.1.1 | $1.90 \times 10^5$<br>$\pm 5.64 \times 10^2$ | $1.28 \times 10^{-3}$<br>$\pm 4.80 \times 10^{-6}$ | 6.674 | 0.151<br>$\pm 1.44 \times 10^{-4}$ | 2.33 x<br>10<sup>-5</sup> |

<sup>a</sup>SE is the standard error for each fitted parameter, <sup>b</sup>SCKAnalyzer calculates the $K_D$ standard error (see also Figures S5, S11 and S17). Table 1 lists only the $K_D$ values obtained from the global analysis of the two replicates. Those obtained by replicate are shown in Figures S4, S10 and S16. <sup>c</sup>Chi<sup>2</sup> is the average of the values calculated by SCKAnalyzer (see Figures S5, S11 and S17).

To further evaluate SCKAnalyzer and how the Octet^®^ R8 BLI instrument performed in SCK mode, SCK4 and SCK5 experiments were conducted. In these new assays an 11-mer DNA duplex was used and the DNA helix stability was tuned through buffer ionic strength and temperature to define experimental conditions for monitoring two distinct binding kinetics. For the SCK5 condition the combination of lower temperature and increased ionic strength relative to SCK4 was expected to enhance DNA helix stability.

The sensorgrams of the SCK4 experiment are shown in Figure S6. For each replicate and blank experiment (buffer-only wells) two sensorgrams were obtained: one for the target-functionalized probe and one without the target. All responses behaved as expected: flat responses were observed with buffer as the analyte or with the probe lacking the target, while clear binding responses occurred when the target-functionalized probe was immersed in wells containing the complementary strand. For each experiment, sensorgrams are saved as .frd files in the results folder generated by Discovery. Consequently, two replicates and one blank experiment (buffer only as analyte) yield three pairs of .frd files. SCKAnalyzer was designed to allow direct upload of these files (Figure S7) and preview the sensorgrams prior to referencing (Figure S8). Once the sensorgrams have been double-referenced and the four injection steps defined they can be fitted. Figure 3A shows the result of the global analysis of the SCK4 experiment using SCKAnalyzer. The corresponding binding parameters are listed in Table 1 (see Figure S9 for the analysis by replicate, and Figure S10 and Figure S11 for the parameters, together with those from the global analysis). As expected for this shorter DNA/DNA duplex compared to the 22-mer RNA/DNA duplex, *K*_D_ is higher, 206 nM. As shown in Figure S12, SCKAnalyzer provides an alternative representation of the fitted sensorgrams by separately overlaying the association and dissociation phases. The fitting results obtained with SCKAnalyzer were then compared to those generated using BIAEvaluation v4.1.1. For this, the sensorgrams from Discovery (BLI acquisition software) (Figure S6) were exported as .dat files (.txt files) and uploaded to BIAevaluation v4.1.1. After double-referencing, the sensorgrams were fitted with a 1:1 model of interaction. The corresponding sensorgrams are shown in Figure 3B and the binding parameters are listed in Table 1 (fits and parameters by replicate are shown in Figure S13, together with the global analysis). As reported in Table 1, both software tools retrieved highly consistent values across all parameters (*k*_a_, *k*_d_, *K*_D_, and *R*_max_).

**Figure 3:**
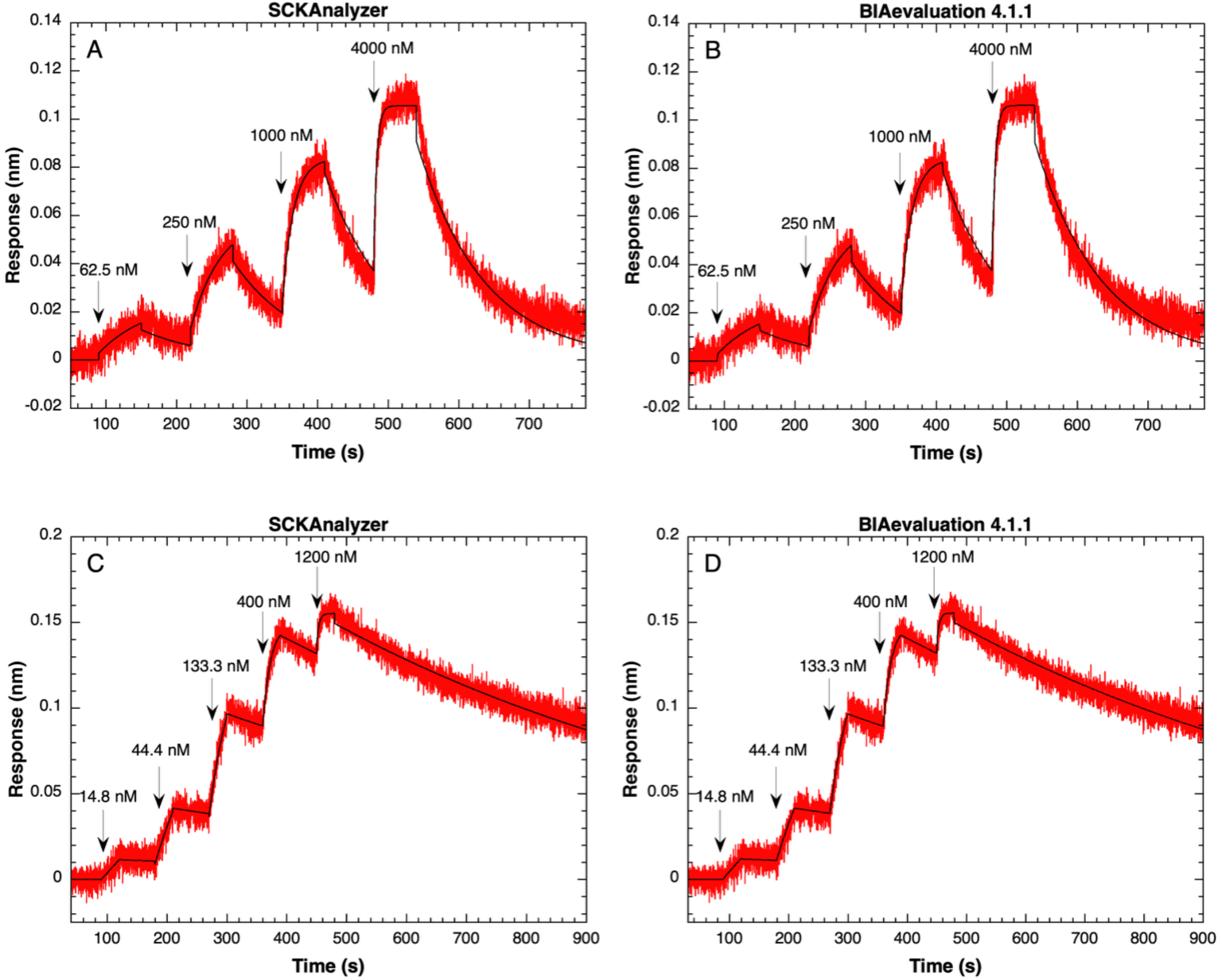
Sensorgrams of the DNA/DNA duplex, analyzed with SCKAnalyzer and BIAevaluation v4.1.1. The red and black lines represent the experimental data and fit to a 1:1 model of interaction, respectively. Arrows indicate injection start times.

To futher assess the performance of SCKAnalyzer on SCK-based BLI data, the same DNA-DNA complex was analyzed using five analyte concentrations, *i.e.* by an SCK5 method, under conditions of decreased temperature and increased ionic strength. *K*_D_ is therefore expected to be lower than 202-206 nM. The sensorgrams of the SCK5 experiments are shown in Figure S14. The corresponding .frd files were uploaded in SCKAnalyzer processed and fitted as described for the SCK4 experiments. The results are shown in Figure 3C and the corresponding parameters are listed in Table 1 (see Figures S15-S17 for the analysis by replicate and the parameters, together with the global analysis, association and dissociation step overlaps are shown in Figure S18). The resulting *K*_D_ was 6.65 nM, consistent with enhanced duplex stability at lower temperature and higher ionic strength. The sensorgrams from Discovery were then exported as .dat files, uploaded in BIAevaluation v4.1.1, processed and fitted as shown in Figure 3D (see Figure S19 for fits and parameters by replicate, together with the global analysis). Once again, both software retrieved nearly identical numerical values for fitted parameters.

## 4. Conclusions

From a methodological perspective, applying SCK to BLI instruments offers a clear trade-off between consumable costs and experimental throughput. While traditional MCK workflows in a parallel format require multiple probes per concentration gradient, for example, twelve probes (six target-functionalized and six reference) for a standard five-concentration assay (not counting the zero concentration) with double-referencing, SCK reduces this footprint to just two probes, one with the target and one without. However, this sensor economy comes at the expense of a longer total runtime. While the combined duration of the regeneration, baseline and association steps is typically around three minutes in total, the dissociation rate (*k*_d_) of the interaction will dictate the required dissociation time, which largely will drive the overall experiment duration. The choice between SCK and MCK therefore depends not only on budgetary constraints regarding sensor consumption, but also on the kinetic properties of the complex under study.

SCKAnalyzer provides a free, robust online platform for processing SCK sensorgram data, demonstrating high accuracy in determining kinetic (*k*_a_, *k*_d_) and affinity (*K*_D_) constants across both SPR (Biacore T200) and BLI (Octet^®^ R8) systems. Benchmarked against established software, Biacore T200 v3.2.1 and BIAevaluation v4.1.1, SCKAnalyzer was validated using model DNA/RNA and DNA/DNA interactions across three distinct kinetic profiles and SCK conditions. The online tool effectively streamlines the adoption of SCK workflows on BLI instruments and any instrument capable of kinetic titration assays, offering a resource-efficient alternative to conventional MCK.

## CRediT authorship contribution statement

**F.M.** coded, designed and evaluated the software, and wrote the manuscript. **J.M.B.** evaluated the software. **T.S.** evaluated the software and critically reviewed the manuscript. **E.R.** managed the web deployment and server infrastructure for the software. **C.D.** conceptualized the study, performed and analyzed the SPR and BLI experiments, designed and evaluated the software, and wrote the manuscript.

## Declaration of generative AI and AI-assisted technologies in the manuscript preparation process

During the preparation of the manuscript, the authors acknowledge the use of Gemini (Google) to correct grammar and improve readability. All outputs were reviewed and edited by the authors who take full responsibility for the final content of the manuscript.

## Acknowledgements

The authors wish to dedicate this manuscript to the memory of Eric Roubin, a great colleague who has always been there to help us solve our server/network problems. His contribution was fundamental in making SCKAnalyzer freely available online.

We thank the Structural Biophysico-Chemistry platform of the European Institute of Chemistry and Biology (IECB, UAR 3033 CNRS / US001 Inserm, Pessac, France) for access to the Biacore T200 and Octet^®^ R8 instruments. Acquisition of the Biacore T200 was supported by the Conseil Régional d’Aquitaine, the GIS-IBiSA and the Cellule Hôtels à Projets of the CNRS. Acquisition of the Octet^®^ R8 was co-funded by the Conseil Régional d’Aquitaine (50%), along with support from academic laboratories (ARNA, IINS, CBMN, POLYMAT, INRAE BFP, IECB, DIPC) and private partners (Novaptech, CovalX Analytics SAS, Surphase). This work was supported by the European Innovation Council (Grant No. 101099652) and the EUSKAMPUS programme (LTC Incubator BioMINT). T.S. would like to thank the Donostia International Physics Center for being hosted on their premises.

## Supplementary data

<u>Supplementary data</u> is available online

## Data availability

Data will be made available on request.

