## Supplementary data for "SCKAnalyzer: A Web Tool for Single-Cycle Kinetics Analysis and Simulation"

[1]Univ. Bordeaux, CNRS, INSERM, ARNA, UMR 5320, U1212, IECB, F-33000 Bordeaux, France.

[2]Univ. Bordeaux, CNRS, INSERM, UAR3033, US001, IECB, F-33000 Bordeaux, France.

†Deceased, August 31, 2026.

[3]POLYMAT and Advanced Polymers and Materials Department, University of the Basque Country, Donostia - San Sebastian, Spain.

[4]Ikerbasque - Basque Foundation for Science, Bilbao, Spain.

[*]Corresponding authors. Univ. Bordeaux, CNRS, INSERM, ARNA, UMR 5320, U1212, IECB, F-33000 Bordeaux, France

1. Experiments performed using the SCK method with three injections of the analyte (SCK3).

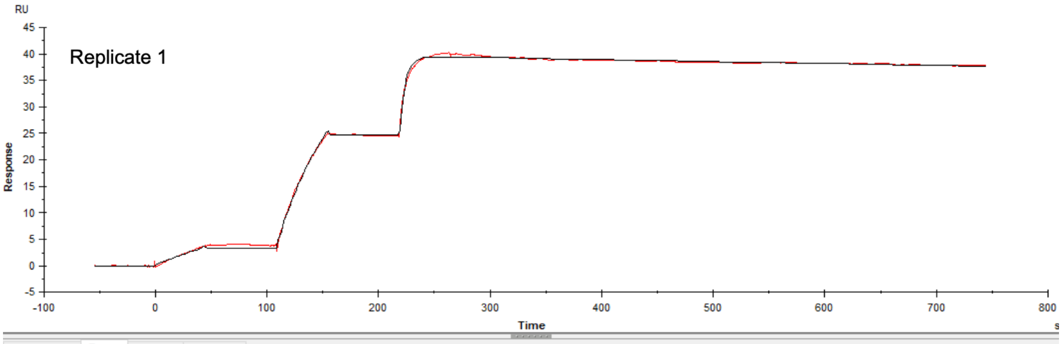


Quality Control | Report | Residuals | Parameters

| Curve | ka (1/Ms) | kd (1/s) | KD (M) | Rmax (RU) | Conc (M) | tc | Flow (ul/min) | kt (RU/Ms) | RI (RU) | Chi² (RU²) | U-value |
|---|---|---|---|---|---|---|---|---|---|---|---|
| | 3.980E+5 | 9.627E-5 | 2.419E-10 | 39.41 | | 1.970E+18 | | | | 0.0645 | 2 |
| Cycle: 11 | | | | | 5.000E-9 | | 25.00 | 5.760E+18 | 0.2507 | | |
| | | | | | 5.000E-8 | | | | 0.6756 | | |
| | | | | | 5.000E-7 | | | | 0.01203 | | |

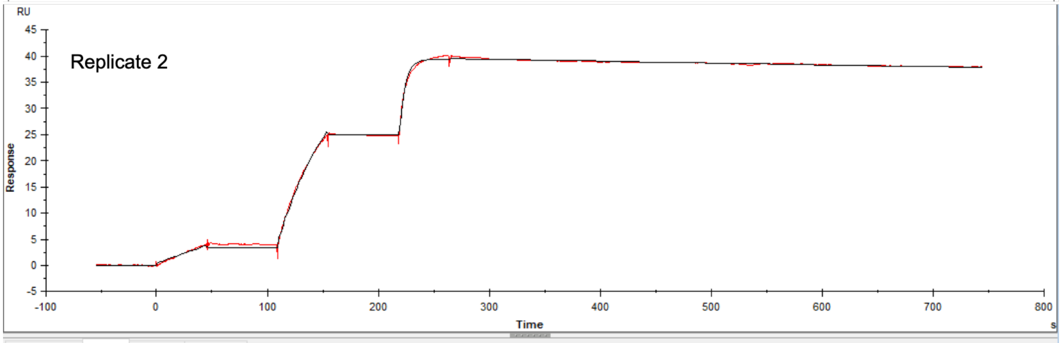


Quality Control | Report | Residuals | Parameters

| Curve | ka (1/Ms) | kd (1/s) | KD (M) | Rmax (RU) | Conc (M) | tc | Flow (ul/min) | kt (RU/Ms) | RI (RU) | Chi² (RU²) | U-value |
|---|---|---|---|---|---|---|---|---|---|---|---|
| | 4.029E+5 | 9.355E-5 | 2.322E-10 | 39.50 | | 6.391E+21 | | | | 0.0722 | 3 |
| Cycle: 12 | | | | | 5.000E-9 | | 25.00 | 1.869E+22 | 0.3863 | | |
| | | | | | 5.000E-8 | | | | 0.8156 | | |
| | | | | | 5.000E-7 | | | | -0.1733 | | |

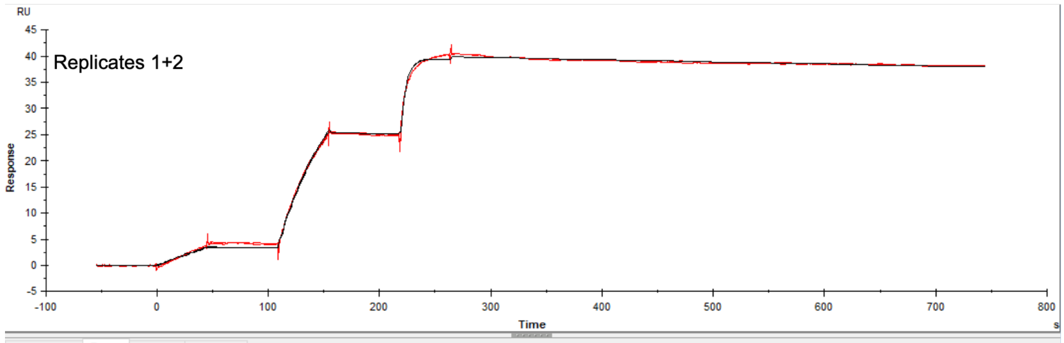


Quality Control | Report | Residuals | Parameters

| Curve | ka (1/Ms) | kd (1/s) | KD (M) | Rmax (RU) | Conc (M) | tc | Flow (ul/min) | kt (RU/Ms) | RI (RU) | Chi² (RU²) | U-value |
|---|---|---|---|---|---|---|---|---|---|---|---|
| | 4.037E+5 | 1.053E-4 | 2.608E-10 | 39.85 | | 1.911E+22 | | | | 0.115 | 3 |
| Cycle: 11 | | | | | 5.000E-9 | | 25.00 | 5.587E+22 | -0.08744 | | |
| | | | | | 5.000E-8 | | | | 0.2284 | | |
| | | | | | 5.000E-7 | | | | -0.4224 | | |
| Cycle: 12 | | | | | 5.000E-9 | | 25.00 | 5.587E+22 | 0.06757 | | |
| | | | | | 5.000E-8 | | | | 0.5453 | | |
| | | | | | 5.000E-7 | | | | -0.4776 | | |

**Figure S1:** Screenshots of SCK3 sensorgrams obtained with the Biacore T200 instrument from the results windows of Biacore T200 Evaluation v3.2.1. The double referenced sensorgrams were fitted to a 1:1: interaction model either individually by replicate (replicate 1, replicate 2) or globally using both replicates simultaneously (replicates 1 + 2 are the sensorgrams shown in Figure 2A with parameters listed in Table 1) to determine the rate constants, $k_a$ and $k_d$, and the dissociation equilibrium constant, $K_D$.

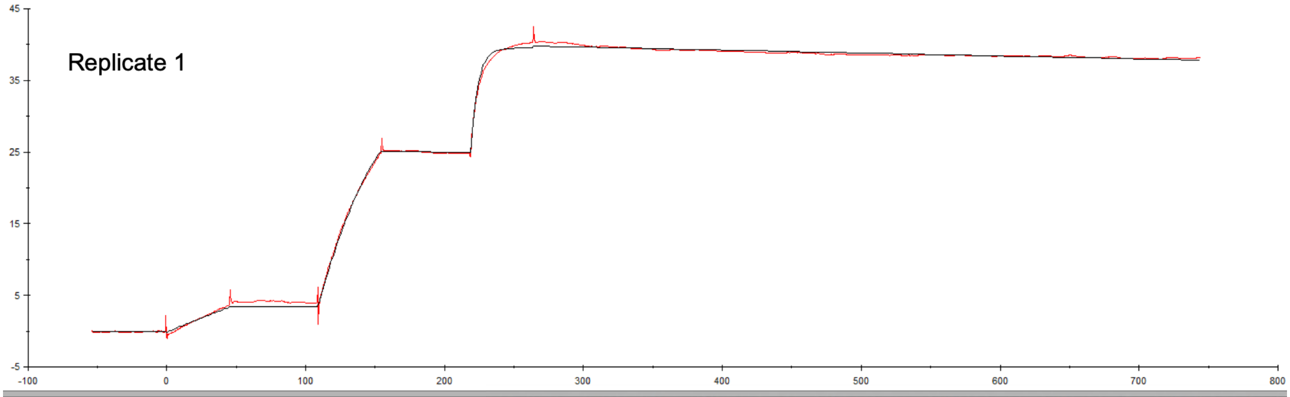


| | R1 | SE(R1) | R2 | SE(R2) | c_time | ton1 | ton2 | F | Conc | Rmax | SE(Rmax) | kt | SE(kt) | ka | SE(ka) | kd | SE(kd) | R3 | SE(R3) | ton3 |
|---|---|---|---|---|---|---|---|---|---|---|---|---|---|---|---|---|---|---|---|---|
| | -0.12 | 0.0156 | 0.0168 | 0.0182 | | | | | | 39.8 | 9.66E-03 | 2.22E+21 | 8.11E+25 | 4.07E+05 | 361 | 1.06E-04 | 8.74E-07 | -0.308 | 0.0177 | |
| Column B vs. Column A | | | | | 45 | 0 | 109 | 10 | 5.00E-07 | | | | | | | | | | | 219 |

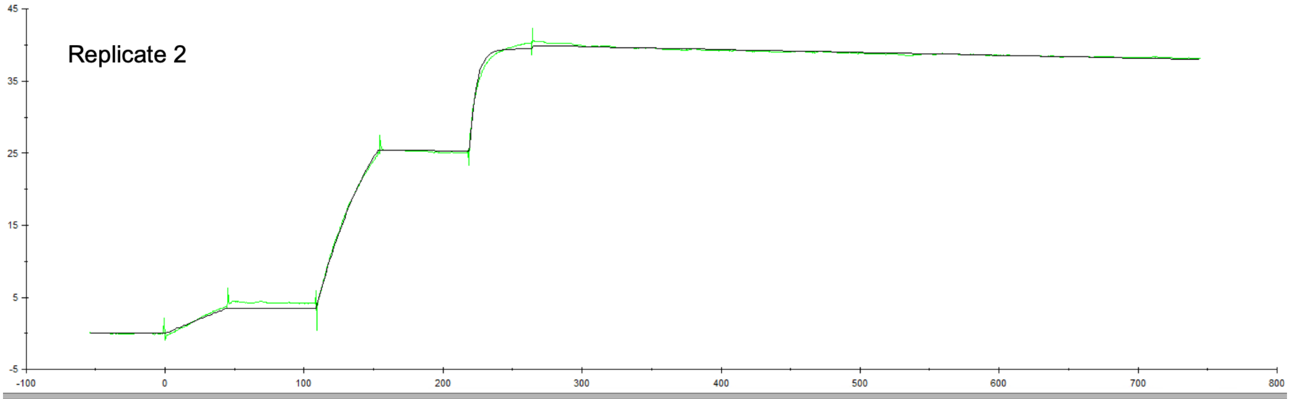


| | R1 | SE(R1) | R2 | SE(R2) | c_time | ton1 | ton2 | F | Conc | Rmax | SE(Rmax) | kt | SE(kt) | ka | SE(ka) | kd | SE(kd) | R3 | SE(R3) | ton3 |
|---|---|---|---|---|---|---|---|---|---|---|---|---|---|---|---|---|---|---|---|---|
| | 9.15E-03 | 0.016 | 0.159 | 0.0186 | | | | | | 39.9 | 9.87E-03 | 6.12E+23 | 1.90E+27 | 4.12E+05 | 373 | 1.03E-04 | 8.91E-07 | -0.492 | 0.0181 | |
| Column F vs. Column E | | | | | 45 | 0 | 109 | 10 | 5.00E-07 | | | | | | | | | | | 219 |

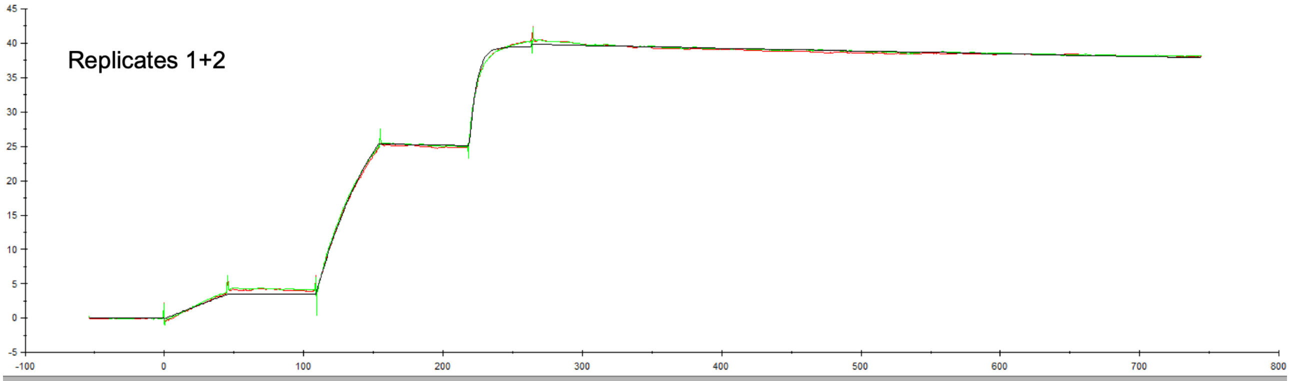


| | R1 | SE(R1) | R2 | SE(R2) | c_time | ton1 | ton2 | F | Conc | Rmax | SE(Rmax) | kt | SE(kt) | ka | SE(ka) | kd | SE(kd) | R3 | SE(R3) | ton3 |
|---|---|---|---|---|---|---|---|---|---|---|---|---|---|---|---|---|---|---|---|---|
| | -0.0552 | 0.0114 | 0.0882 | 0.0133 | | | | | | 39.8 | 7.06E-03 | 2.78E+22 | 1.17E+27 | 4.10E+05 | 265 | 1.05E-04 | 6.38E-07 | -0.4 | 0.0129 | |
| Column B vs. Column A | | | | | 45 | 0 | 109 | 10 | 5.00E-07 | | | | | | | | | | | 219 |
| Column F vs. Column E | | | | | 45 | 0 | 109 | 10 | 5.00E-07 | | | | | | | | | | | 219 |

**Figure S2:** Screenshots of SCK3 sensorgrams obtained with the Biacore T200 instrument from the results windows of BIAevaluation v4.1.1. The sensorgrams from the previous results windows (Figure S1) were exported as .txt files, uploaded in BIAevaluation v4.1.1 and fitted to a 1:1 interaction model either individually by replicate (replicate 1, replicate 2) or globally using both replicates simultaneously (replicates 1 + 2 are the sensorgrams shown in Figure 2B with parameters listed in Table 1) to determine the rate constants, $k_a$ and $k_d$, and the dissociation equilibrium constant, $K_D$.

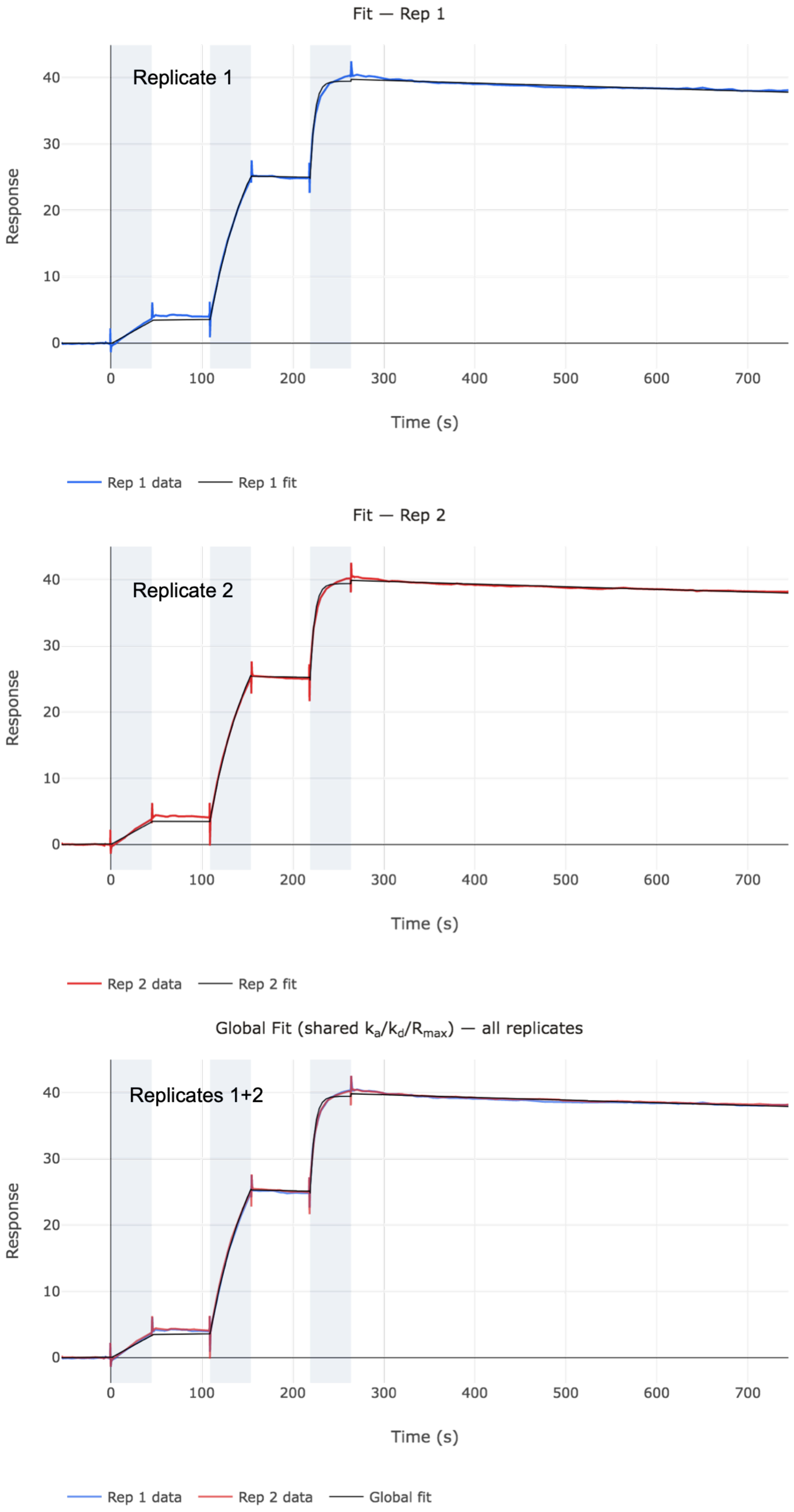


**Figure S3:** Screenshots of SCK3 sensorgrams obtained with the Biacore T200 instrument from the results windows of SCKAnalyzer. The sensorgrams from the previous results windows (Figure S1) were exported as .txt files, uploaded in SCKAnalyzer and merged as a new file the Prepare data set menu of SCKAnalyzer. The sensorgrams were fitted to a 1:1 interaction model either individually by replicate (Replicate 1, Replicate 2) or globally using both replicates simultaneously (Replicates 1 + 2 are the sensorgrams shown in Figure 2C with parameters listed in Table 1) to determine the rate constants, $k_a$ and $k_d$, and the dissociation equilibrium constant, $K_D$.

Fit results

| | $k_a$ (1/M·s) | SE | $k_d$ (1/s) | SE | $K_D$ (M) | SE | $R_{max}$ (RU) | SE |
|---|---|---|---|---|---|---|---|---|
| Rep 1 | 4.0662e+5 | 3.793e+2 | 1.0429e-4 | 9.187e-7 | 2.5649e-10 | 2.272e-12 | 3.9774e+1 | 1.029e-2 |
| Rep 2 | 4.1133e+5 | 3.925e+2 | 1.0225e-4 | 9.213e-7 | 2.4857e-10 | 2.252e-12 | 3.9877e+1 | 1.030e-2 |

Summary (2 replicates)

| Parameter | Mean | SD | Unit |
|---|---|---|---|
| $k_a$ | 4.0897e+5 | 3.3346e+3 | 1/M·s |
| $k_d$ | 1.0327e-4 | 1.4461e-6 | 1/s |
| $K_D$ | 2.5253e-10 | 5.5950e-12 | M |
| $R_{max}$ | 3.9826e+1 | 7.2577e-2 | RU |

Fit quality

| | RMSE (RU) | $R^2$ | $Chi^2$ | N points |
|---|---|---|---|---|
| Rep 1 | 0.3297 | 0.9995 | 1.0878e-1 | 7990 |
| Rep 2 | 0.3371 | 0.9995 | 1.1370e-1 | 7990 |

Bulk offsets

| | Inj 1 (RU) | Inj 2 (RU) | Inj 3 (RU) |
|---|---|---|---|
| Rep 1 | -0.111 | 0.052 | -0.274 |
| Rep 2 | 0.018 | 0.202 | -0.471 |

**Figure S4:** Screenshot showing the SCKAnalyzer fit results window for the analysis by replicate.

Fit results

| | $k_a$ (1/M·s) | SE | $k_d$ (1/s) | SE | $K_D$ (M) | SE | $R_{max}$ (RU) | SE |
|---|---|---|---|---|---|---|---|---|
| Global fit | 4.0899e+5 | 2.808e+2 | 1.0336e-4 | 6.659e-7 | 2.5272e-10 | 1.637e-12 | 3.9827e+1 | 7.452e-3 |

Fit quality

| | RMSE (RU) | $R^2$ | $Chi^2$ | N points |
|---|---|---|---|---|
| Rep 1 | 0.3367 | 0.9995 | 1.1344e-1 | 7990 |
| Rep 2 | 0.3452 | 0.9995 | 1.1923e-1 | 7990 |

Bulk offsets

| | Inj 1 (RU) | Inj 2 (RU) | Inj 3 (RU) |
|---|---|---|---|
| Rep 1 | -0.048 | 0.125 | -0.375 |
| Rep 2 | -0.048 | 0.125 | -0.375 |

**Figure S5:** Screenshot showing the SCKAnalyzer fit results window for the global analysis (parameters listed in Table 1).

2. Experiments performed using the SCK method with four injections of the analyte (SCK4).

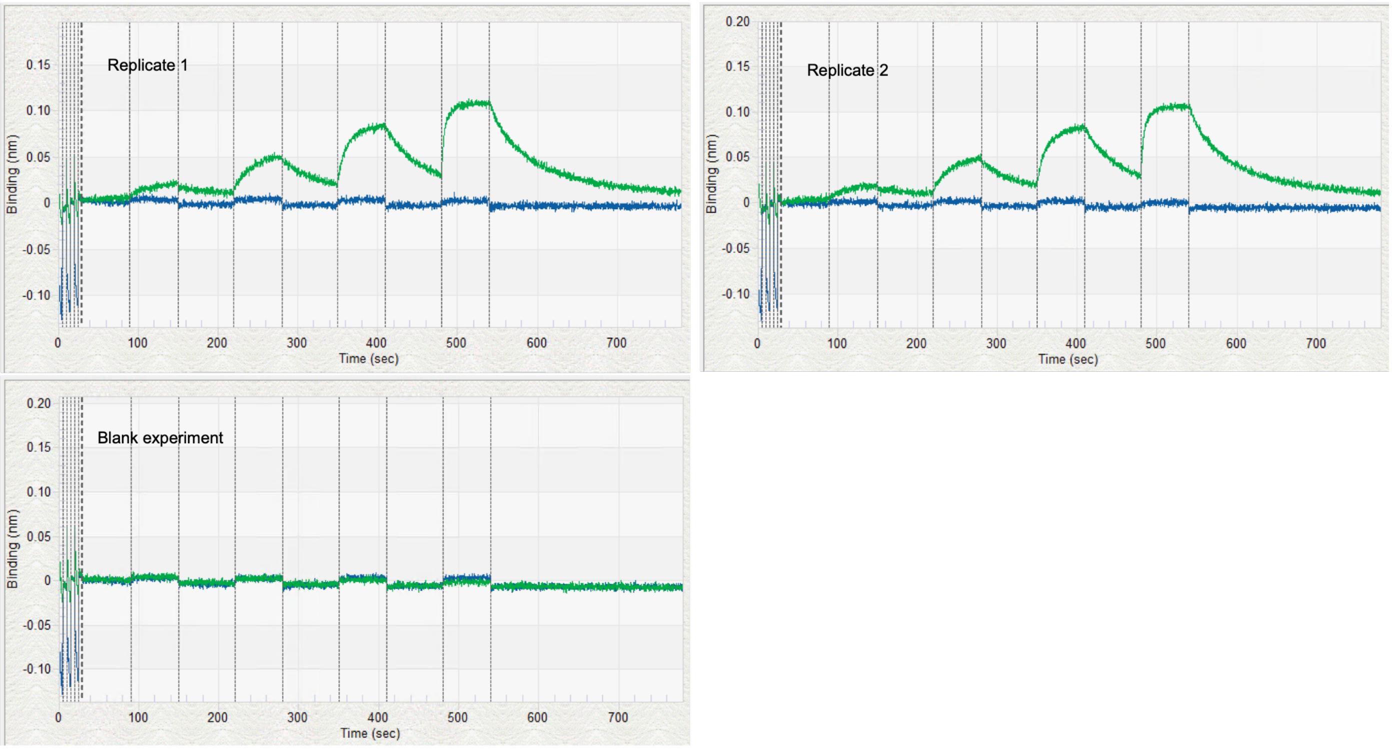


**Figure S6:** Screenshots of the experiments performed on the Octet R8 instrument monitored with the Discovery v13.1.1.29 acquisition software. One probe was left blank (blue lines) and the other one was functionalized with the biotinylated target (green lines, see Materials and Methods). The analyte was prepared in different wells on the x-axis of the 96-well plate at increasing concentrations of 62.5, 250, 1000, and 4000 nM (Replicate1, Replicate2). Both probes, with and without target, were immersed for 60 s in the same analyte samples during the titration. One experiment was performed with running buffer as analyte for double-referencing of the sensorgrams (Blank experiment). For each experiment, the Discovery acquisition software saves the data as two .frd files, one for the probe without the target and one for the probe with the target, resulting in three pairs of .frd files total for two replicates plus a blank experiment.

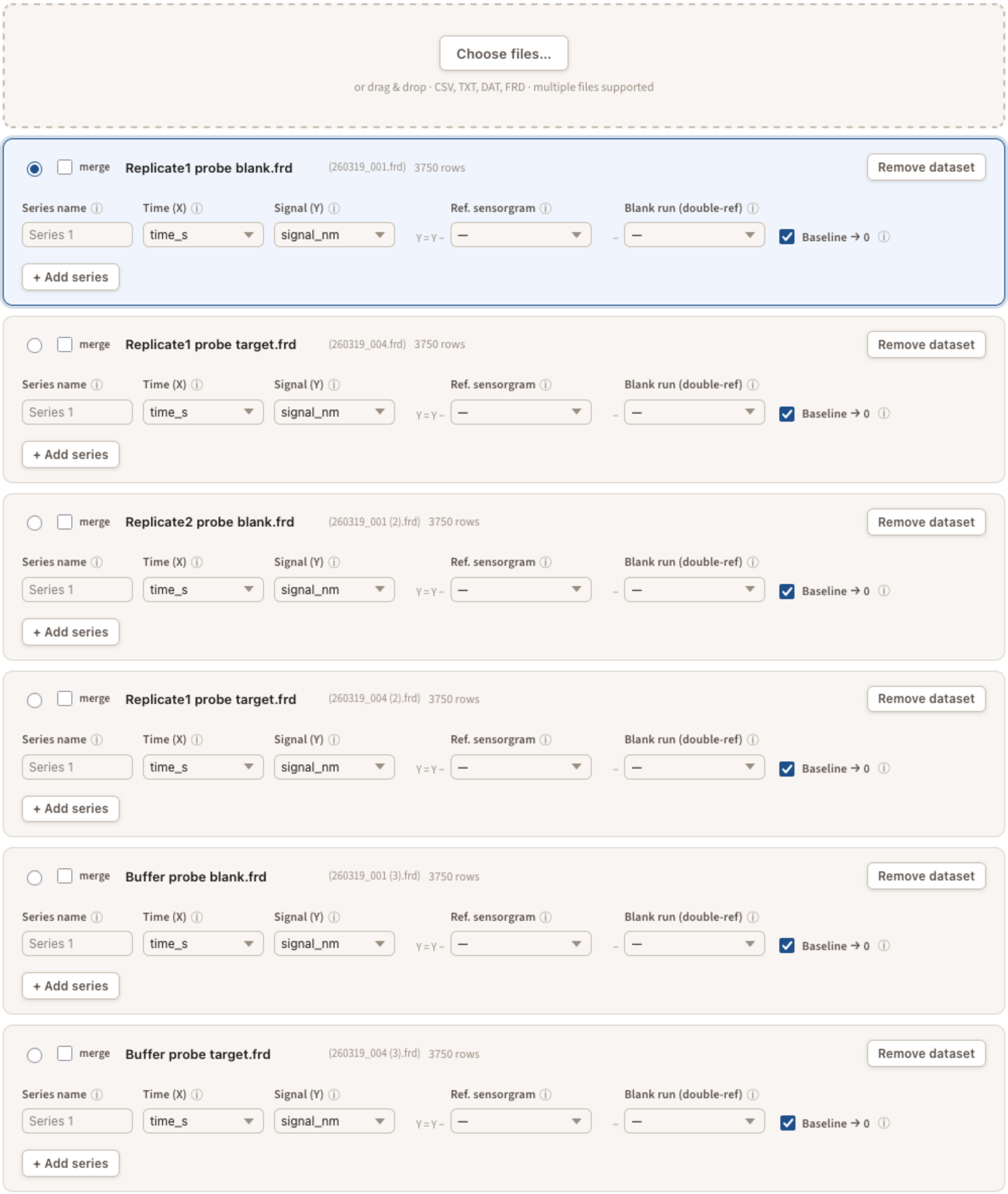


**Figure S7:** Screenshot of the Prepare dataset menu of SCKAnalyzer for the SCK4 experiment. The data save as .frd files in BLI Discovery acquisition software were uploaded in SCKAnalyzer and rename per replicate plus the experiment with buffer as analyte. In the file names, “Replicate 1” and “Replicate 2” indicate assays performed with the analyte, while “Buffer” indicates the control experiment performed with buffer as the analyte." “Probe target” and “Probe blank” in the name file indicate probes with and without the biotinylated target, respectively. The referenced sensorgrams were obtained by subtracting the signal of the “probe blank” from the “probe target” in the “Ref. sensorgram” menu. The double-referenced sensorgrams were obtained by then substracting the referenced buffer file from the referenced replicates in “Blank run (double-ref)”.

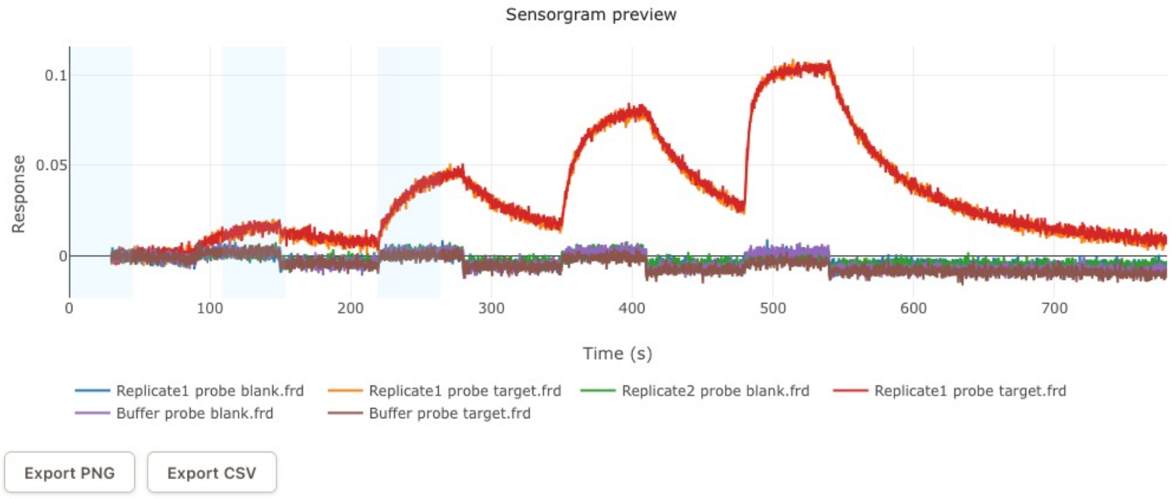


**Figure S8:** Screenshot of the Sensorgram preview. The sensorgrams correspond to those uploaded in the Prepare dataset (Figure S7). Each experiment (Replicate1, Replicate2, Buffer) has two .frd files, one for the probe with the target and one without.

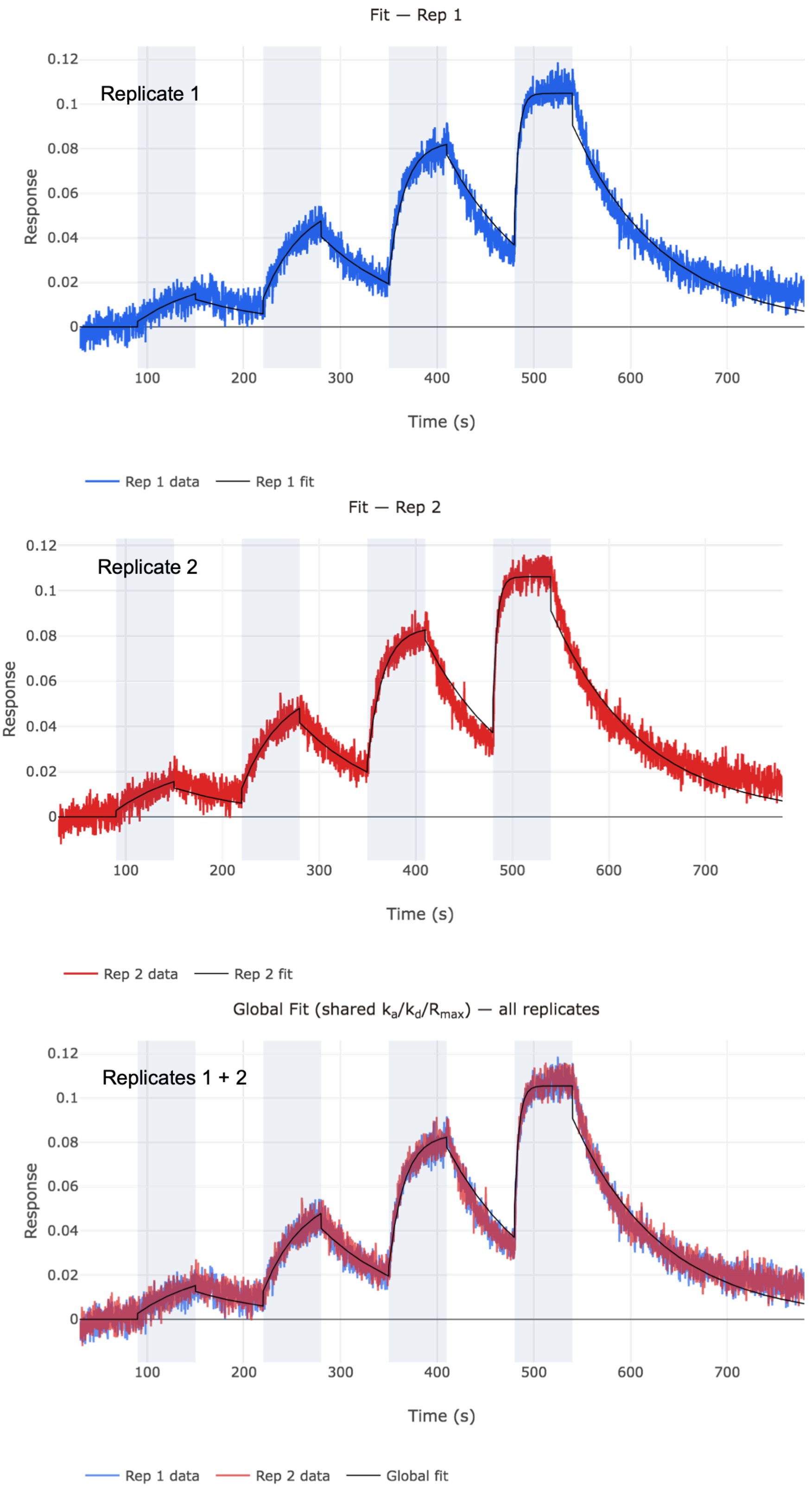


**Figure S9:** Screenshots of SCK4 sensorgrams obtained with the Octet R8 instrument from the results windows of SCKAnalyzer. The sensorgrams from the BLI assay (Figure S6) were uploaded in SCKAnalyzer directly as .frd files and fitted to a 1:1 interaction model either individually by replicate (Replicate 1, Replicate 2) or globally using both replicates simultaneously (Replicates1 + Rep 2 Global Fit are sensorgrams shown in Figure 3A with parameters listed in Table 1) to determine the rate constants, $k_a$ and $k_d$, and the dissociation equilibrium constant, $K_D$.

Fit results

| | $k_a$ (1/M·s) | SE | $k_d$ (1/s) | SE | $K_D$ (M) | SE | $R_{max}$ (RU) | SE |
|---|---|---|---|---|---|---|---|---|
| Rep 1 | 5.0897e+4 | 6.733e+2 | 1.0639e-2 | 7.831e-5 | 2.0902e-7 | 3.164e-9 | 9.5157e-2 | 4.733e-4 |
| Rep 2 | 5.2487e+4 | 7.082e+2 | 1.0654e-2 | 8.013e-5 | 2.0299e-7 | 3.136e-9 | 9.5735e-2 | 4.839e-4 |

Summary (2 replicates)

| Parameter | Mean | SD | Unit |
|---|---|---|---|
| $k_a$ | 5.1692e+4 | 1.1241e+3 | 1/M·s |
| $k_d$ | 1.0646e-2 | 1.0973e-5 | 1/s |
| $K_D$ | 2.0600e-7 | 4.2676e-9 | M |
| $R_{max}$ | 9.5446e-2 | 4.0859e-4 | RU |

Fit quality

| | RMSE (RU) | $R^2$ | $Chi^2$ | N points |
|---|---|---|---|---|
| Rep 1 | 0.0053 | 0.9681 | 2.8341e-5 | 3750 |
| Rep 2 | 0.0055 | 0.9667 | 3.0101e-5 | 3750 |

Bulk offsets

| | Inj 1 (RU) | Inj 2 (RU) | Inj 3 (RU) | Inj 4 (RU) |
|---|---|---|---|---|
| Rep 1 | 0.003 | 0.007 | 0.005 | 0.014 |
| Rep 2 | 0.003 | 0.007 | 0.004 | 0.015 |

**Figure S10:** Screenshot showing the SCKAnalyzer fit results window for the analysis by replicate.

Fit results

| | $k_a$ (1/M·s) | SE | $k_d$ (1/s) | SE | $K_D$ (M) | SE | $R_{max}$ (RU) | SE |
|---|---|---|---|---|---|---|---|---|
| Global fit | 5.1674e+4 | 4.889e+2 | 1.0647e-2 | 5.610e-5 | 2.0604e-7 | 2.231e-9 | 9.5451e-2 | 3.390e-4 |

Fit quality

| | RMSE (RU) | $R^2$ | $Chi^2$ | N points |
|---|---|---|---|---|
| Rep 1 | 0.0053 | 0.9680 | 2.8446e-5 | 3750 |
| Rep 2 | 0.0055 | 0.9666 | 3.0206e-5 | 3750 |

Bulk offsets

| | Inj 1 (RU) | Inj 2 (RU) | Inj 3 (RU) | Inj 4 (RU) |
|---|---|---|---|---|
| Rep 1 | 0.003 | 0.007 | 0.005 | 0.015 |
| Rep 2 | 0.003 | 0.007 | 0.005 | 0.015 |

**Figure S11:** Screenshot showing the SCKAnalyzer fit results window for the global analysis (parameters listed in Table 1).

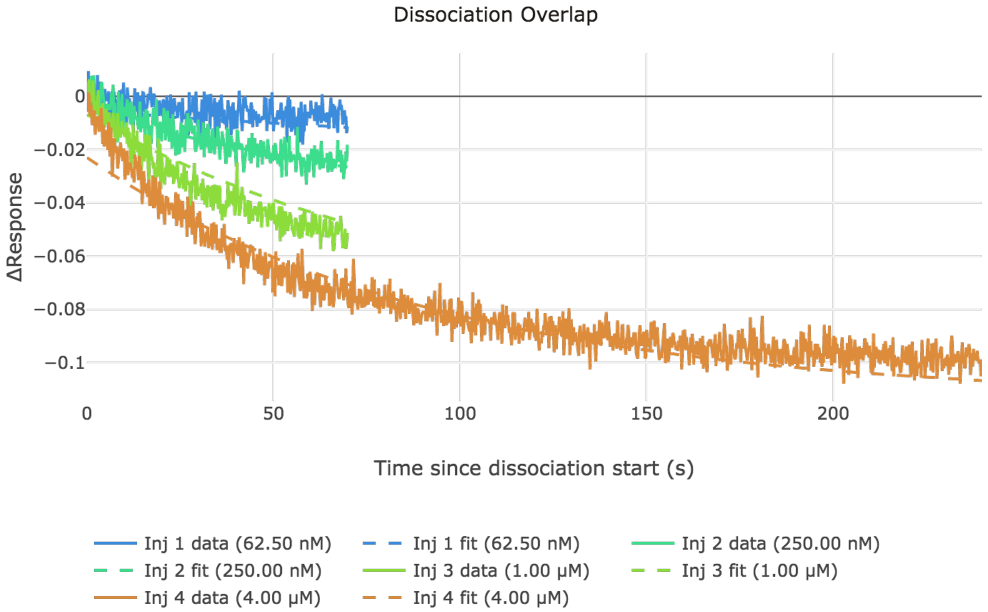


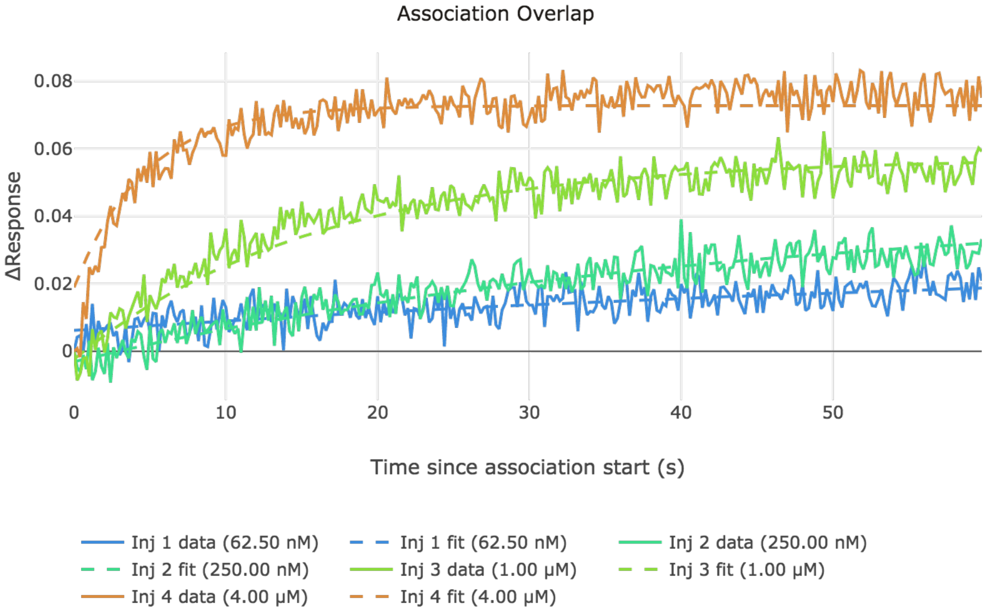


**Figure S12**: Response and fit for each association and dissociation phases. The two figures correspond to the first replicate of the global fit analysis.

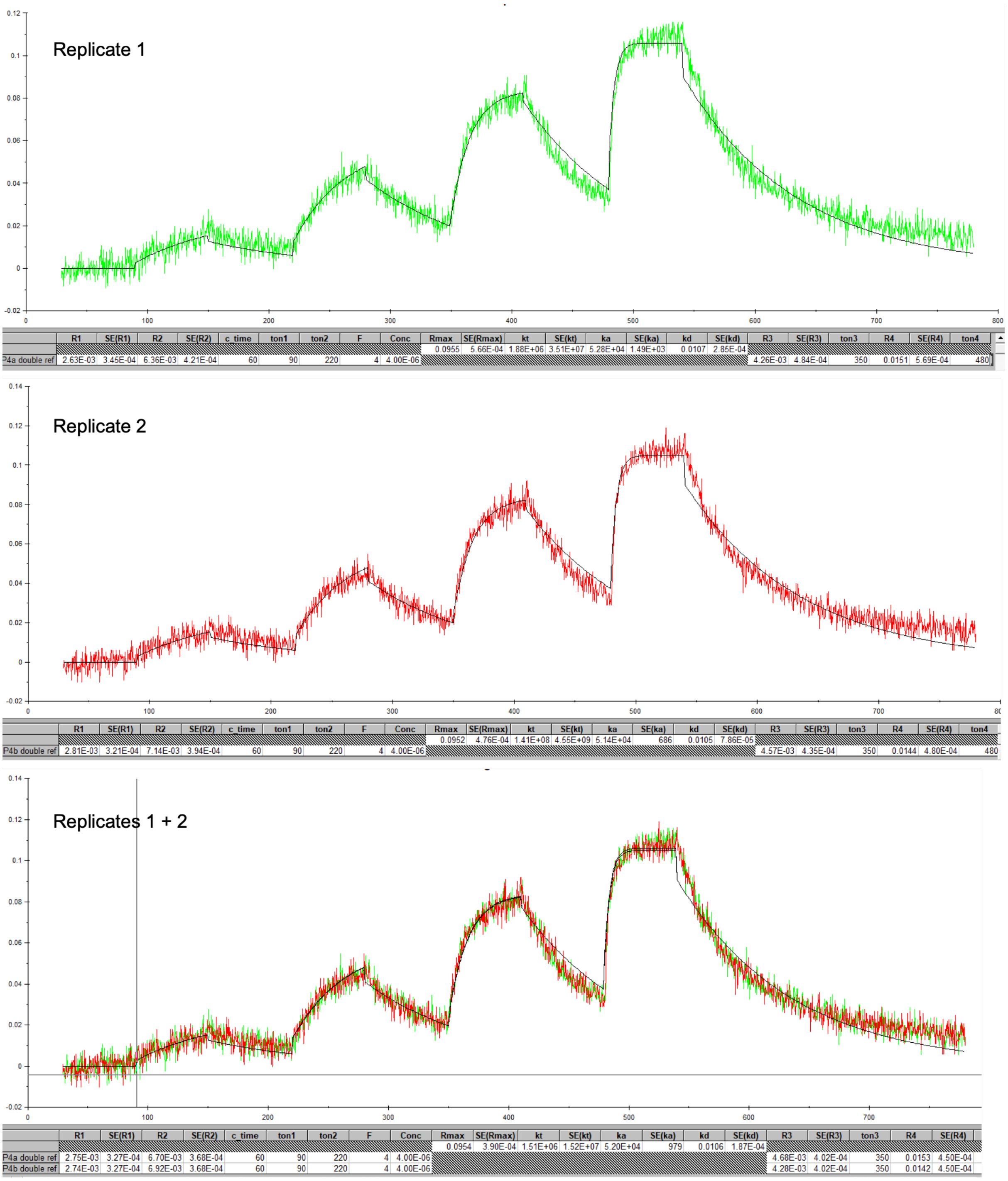


**Figure S13:** Screenshots of SCK4 sensorgrams obtained with the Octet R8 instrument from the results windows of BIAevaluation v4.1.1. The sensorgrams shown in Figure S6 were exported from Discovery windows (Figure S6) as .dat files and uploaded in BIAevaluation v4.1.1. After double referencing they were fitted to a 1:1 interaction model either individually by replicate (replicate 1, replicate 2) or globally using both replicates simultaneously (replicates 1 + 2 are the sensorgrams shown in Figure 3B with parameters listed in Table 1) to determine the rate constants, $k_a$ and $k_d$, and the dissociation equilibrium constant, $K_D$.

3. Experiments performed using the SCK method with five injections of the analyte (SCK5).

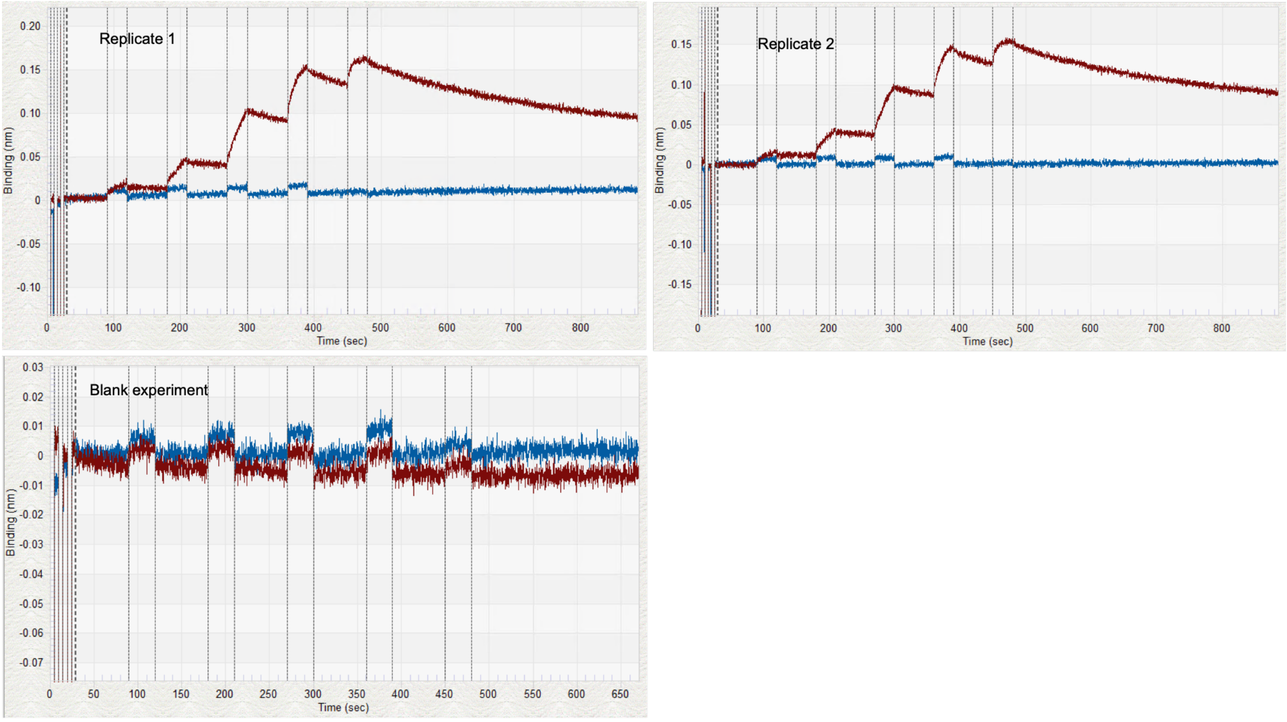


**Figure S14:** Screenshots of the experiments performed on the Octet R8 instrument monitored with the Discovery v13.1.1.29 acquisition software. One probe was left blank and the other one was functionalized with the biotinylated target (see Materials and Methods). The analyte was injected at increasing concentrations of 14.8, 44.4, 133.3, 400, and 1200 nM for 30 s (Replicate1, Replicate2). One experiment was performed with running buffer as analyte for double-referencing of the sensorgrams.

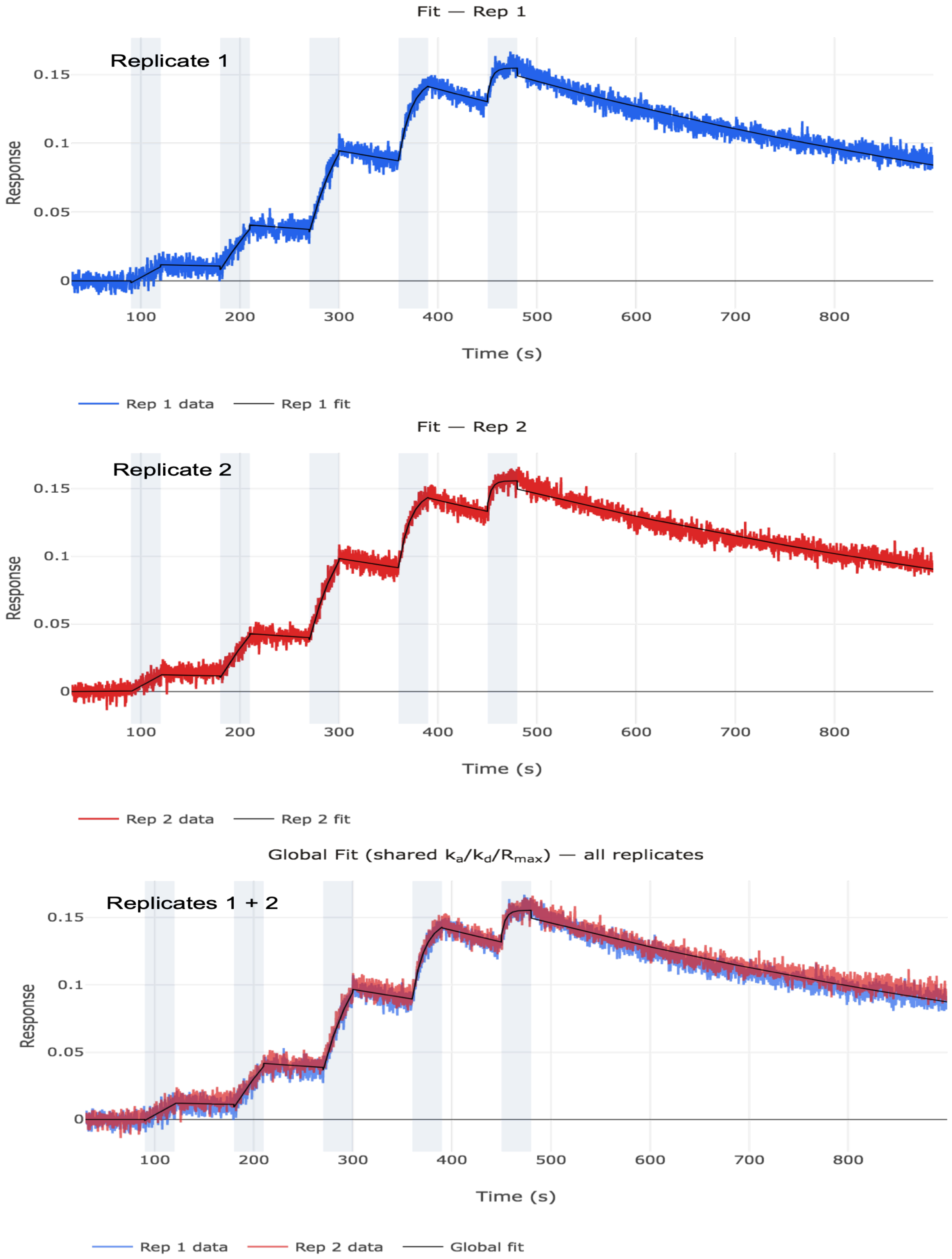


**Figure S15:** Screenshots of SCK5 sensorgrams obtained with the Octet R8 instrument from the results windows of SCKAnalyzer. The sensorgrams from the BLI assay (Figure S15) were uploaded in SCKAnalyzer directly as *.frd* files and fitted to a 1:1 interaction model either individually by replicate (Rep 1, Rep 2) or globally using both replicates simultaneously (Rep 1 + Rep 2 Global Fit are the sensorgrams shown in Figure 3C with parameters listed in Table 1) to determine the rate constants, $k_a$ and $k_d$, and the dissociation equilibrium constant, $K_D$.

### Fit results

| | $k_a$ (1/M·s) | SE | $k_d$ (1/s) | SE | $K_D$ (M) | SE | $R_{max}$ (RU) | SE |
|---|---|---|---|---|---|---|---|---|
| Rep 1 | 1.8729e+5 | 8.378e+2 | 1.3622e-3 | 6.713e-6 | 7.2736e-9 | 4.841e-11 | 1.5020e-1 | 2.039e-4 |
| Rep 2 | 1.9772e+5 | 8.767e+2 | 1.1998e-3 | 6.464e-6 | 6.0684e-9 | 4.234e-11 | 1.5046e-1 | 1.994e-4 |

Summary (2 replicates)

| Parameter | Mean | SD | Unit |
|---|---|---|---|
| $k_a$ | 1.9250e+5 | 7.3752e+3 | 1/M·s |
| $k_d$ | 1.2810e-3 | 1.1485e-4 | 1/s |
| $K_D$ | 6.6710e-9 | 8.5219e-10 | M |
| $R_{max}$ | 1.5033e-1 | 1.8141e-4 | RU |

Fit quality

| | RMSE (RU) | $R^2$ | $Chi^2$ | N points |
|---|---|---|---|---|
| Rep 1 | 0.0045 | 0.9916 | 2.0040e-5 | 4350 |
| Rep 2 | 0.0045 | 0.9916 | 2.0276e-5 | 4350 |

Bulk offsets

| | Inj 1 (RU) | Inj 2 (RU) | Inj 3 (RU) | Inj 4 (RU) | Inj 5 (RU) |
|---|---|---|---|---|---|
| Rep 1 | -0.001 | -0.003 | -0.002 | 0.000 | 0.006 |
| Rep 2 | 0.000 | -0.001 | -0.002 | 0.001 | 0.006 |

**Figure S16:** Screenshot showing the SCKAnalyzer fit results window for the analysis by replicate.

### Fit results

| | $k_a$ (1/M·s) | SE | $k_d$ (1/s) | SE | $K_D$ (M) | SE | $R_{max}$ (RU) | SE |
|---|---|---|---|---|---|---|---|---|
| Global fit | 1.9239e+5 | 6.485e+2 | 1.2797e-3 | 4.984e-6 | 6.6514e-9 | 3.426e-11 | 1.5032e-1 | 1.526e-4 |

Fit quality

| | RMSE (RU) | $R^2$ | $Chi^2$ | N points |
|---|---|---|---|---|
| Rep 1 | 0.0048 | 0.9904 | 2.2992e-5 | 4350 |
| Rep 2 | 0.0048 | 0.9903 | 2.3234e-5 | 4350 |

Bulk offsets

| | Inj 1 (RU) | Inj 2 (RU) | Inj 3 (RU) | Inj 4 (RU) | Inj 5 (RU) |
|---|---|---|---|---|---|
| Rep 1 | -0.001 | -0.002 | -0.002 | 0.001 | 0.006 |
| Rep 2 | -0.001 | -0.002 | -0.002 | 0.001 | 0.006 |

**Figure S17:** Screenshot showing the SCKAnalyzer fit results window for the global analysis (parameter listing in Table 1).

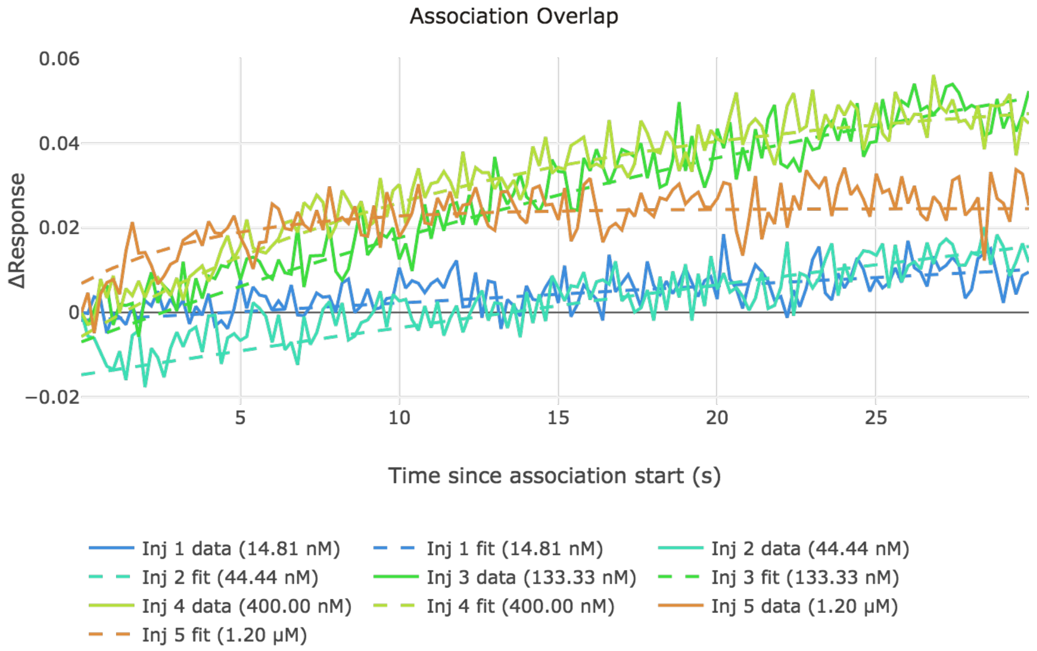


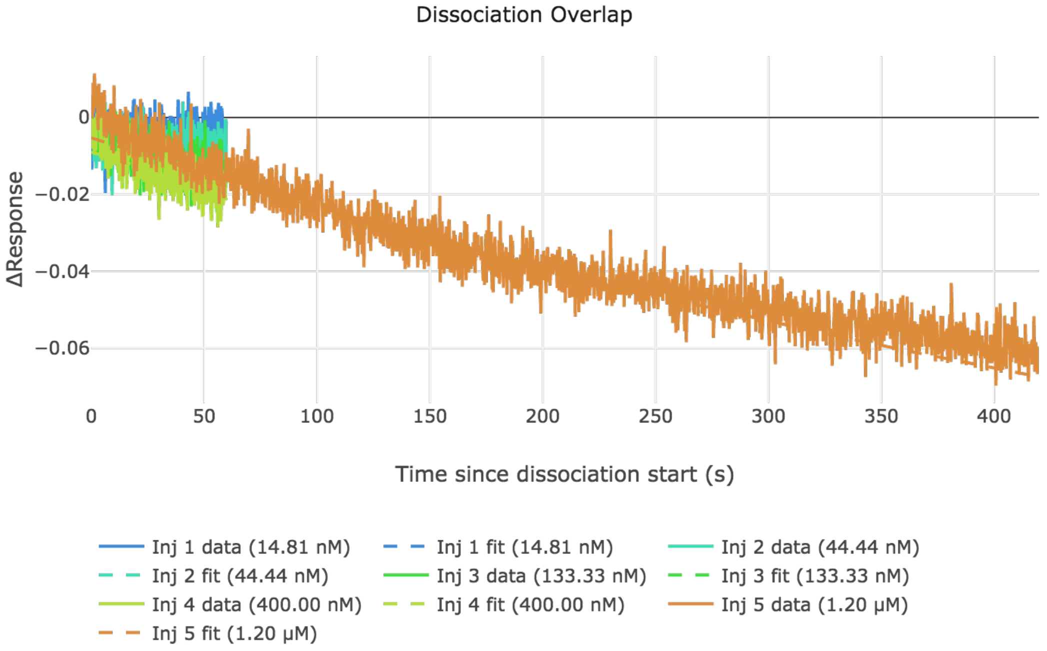


**Figure S18**: Response and fit for each association and dissociation phases. The two figures correspond to the first replicate of the global fit analysis.

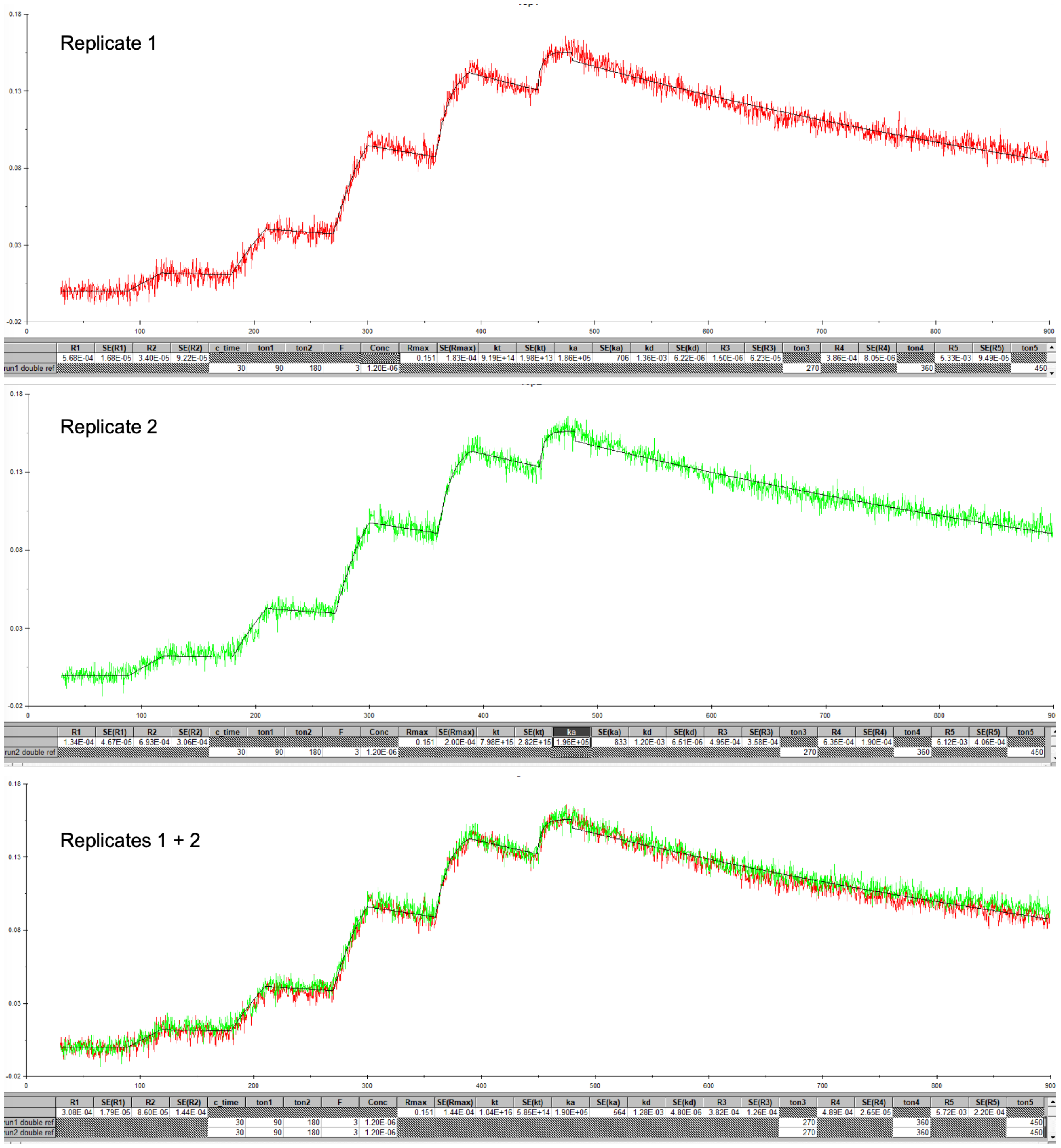


**Figure S19:** Screenshots of SCK5 sensorgrams obtained with the Octet R8 instrument from the results windows of BIAevaluation v4.1.1. The sensorgrams shown in Figure S6 were exported from Discovery windows (Figure S14) as .dat files and uploaded in BIAevaluation v4.1.1. After double-referencing they were fitted to a 1:1 interaction model either individually by replicate (replicate 1, replicate 2) or globally using both replicates simultaneously (replicates 1 + 2 are the sensorgrams shown in Figure 3D with parameter listed in Table 1) to determine the rate constants, $k_a$ and $k_d$, and the dissociation equilibrium constant, $K_D$.
